# Impact of sequencing approach on ecological inference from highly variable bat gut microbiomes

**DOI:** 10.64898/2026.01.08.698297

**Authors:** Ostaizka Aizpurua, Evie Morris, Lasse Nyholm, Raphael Eisenhofer, Carmi Korine, Amy Gladwell, Orly Razgour, Antton Alberdi

## Abstract

Understanding the functional potential of animal gut microbiomes requires analytical approaches that can accurately capture both taxonomic and functional diversity, particularly in hosts with samples with low and variable microbial DNA fractions, such as bats. Here, we compared 16S amplicon sequencing and genome-resolved metagenomics for characterising the gut microbiomes of three sympatric insectivorous bats (*Pipistrellus kuhlii*, *Hypsugo ariel*, and *Cnephaeus bottae*). Amplicon sequencing recovered 3,536 amplicon sequence variants (ASVs) spanning 29 microbial phyla, including five archaeal groups, while metagenomics yielded 135 metagenome-assembled genomes (MAGs) across 15 bacterial phyla. Alpha and beta diversity patterns differed significantly between approaches. Applying prevalence- and abundance-based filtering to amplicon data improved alpha diversity congruence with metagenomic profiles, indicating that low-abundance and rare ASVs inflate diversity estimates, while the low microbial DNA fractions in bat faeces challenge genome reconstruction of such taxa. Functional reconstructions diverged notably, with metagenomics-based functional profiles yielding consistently lower metabolic capacities than those indirectly inferred from 16S amplicon sequencing, both before and after prevalence- and abundance-based filtering, due to incompleteness of many reconstructed genomes. However, metagenome-assembled genomes harboured metabolic capacities missed by the indirect functional inference of metabarcoding, which largely rely on genomes of strains characterised in model organisms. Together, these findings highlight the influence of chosen methodology, host species, and data filtering on microbiome profiling in taxa with low microbial biomass. We propose practical guidelines to improve analytical consistency and reliability when studying gut microbiomes of bats and other hosts with similarly variable microbial communities.

## Introduction

Many biological processes in animals are strongly influenced by the microorganisms associated with them (McFall-Ngai et al., 2013). Commensal members of the gut microbiome serve not only as a defensive shield against pathogens (McLaren & Callahan, 2020) but also play pivotal roles in regulating nutrient absorption and energy balance, among other critical functions (Flint et al., 2012). Gut microbiomes vary markedly with diet, exposure to pollutants, and other ecological factors, conferring a plasticity that may facilitate adaptation to new environments (Alberdi et al., 2016). Consequently, there is a growing acknowledgement of the need to factor in gut microbiomes when studying animal biology (Carthey et al., 2020).

Presently, two primary methodological approaches are employed to analyse gut microbiomes. Amplicon sequencing of the 16S rRNA gene (hereafter, metabarcoding) is the most popular approach due to its simplicity and reduced cost (Poretsky et al., 2014). Through this strategy, researchers analyse the sequences of a PCR-amplified standardised genetic marker to infer the taxonomic composition and relative abundances of bacteria within samples (Prodan et al., 2020). Functional attributes of taxa and communities can then be indirectly inferred from genome databases by matching the profiled 16S rRNA sequence with the functional attributes of reference genomes with similar 16S rRNA gene (Iwai et al., 2016; Langille et al., 2013). In contrast, genome-resolved shotgun metagenomics (hereafter, metagenomics) relies on the sequencing of total DNA present in the samples (Quince et al., 2017). Complex bioinformatic procedures are then applied to reconstruct metagenome-assembled genomes (MAGs), which enable profiling of taxonomic and functional features of microbial communities. Notably, metagenomics offers a distinct advantage over metabarcoding through the direct capture of functional traits from the reconstructed genomes (Koziol et al., 2023), theoretically providing a more accurate representation of the functional attributes of microbiomes (Eloe-Fadrosh et al., 2016; Liu et al., 2020).

Recent research on vertebrate-associated microbial communities has showcased the power of genome-resolved metagenomics in characterising the functional attributes of wild microbiomes (Leonard et al., 2024; Levin et al., 2021; Worsley et al., 2024). However, host biological traits can influence the recovery of microbial genomes and the overall performance of microbiome profiling (Aizpurua et al., 2023). A large comparative study spanning over 150 species found that genome-resolved metagenomics produces consistent results in non-flying mammals, reptiles, and amphibians, but outcomes are notably more variable in birds and bats (Pietroni et al., 2024). Human and mouse faecal samples, which are among the most extensively studied (Arumugam et al., 2011; Laukens et al., 2016), tend to exhibit relatively stable and predictable microbial biomass and composition (Tang et al., 2025). In contrast, samples from bats and birds typically contain a lower proportion of bacterial and archaeal DNA and display greater variability, resulting in pronounced fluctuations in measured microbiome diversity (Pietroni et al., 2024). These disparities present challenges for identifying the most accurate analytical approach, as data generated under current community standards may be insufficient for reliable interpretation (Aizpurua et al., 2023).

To evaluate the strengths and limitations of different approaches for resolving functional features in challenging microbiome samples, we compared metabarcoding- and metagenomics-based strategies for characterising both the taxonomic and functional attributes of gut microbiomes in three sympatric insectivorous bats: *Pipistrellus kuhlii*, *Hypsugo ariel*, and *Cnephaeus* (formerly *Eptesicus*) *bottae (Cláudio et al., 2023)*. Based on knowledge of the ecology of the three bat species (Amichai & Korine, 2020; Korine, 2021; Yom-tov & Kadmon, 1998), we compared three contrasting hypotheses: 1) Shared diet hypothesis - *P. kuhlii* and *H. ariel* share more similar gut microbiome composition, diversity and functionality than *C. bottae* due to their similar size and diet (Feldman et al., 2000). 2) Shared habitat hypothesis - *P. kuhlii* and *C. bottae* share more similar gut microbiome than *H. ariel* because they are found in both natural desert and anthropogenic habitats, unlike *H. ariel* that is mostly found in hyper-arid natural desert habitats (Korine & Pinshow, 2004). 3) Shared evolutionary history hypothesis - *H. ariel* and *C. bottae* share more similar gut microbiome because they are restricted to desert environments, while *P. kuhlii* is a Mediterranean species that has recently expanded its distribution to desert environments following the spread of urban areas and irrigated agriculture (Yom-Tov & Tchernov, 1988).

Specifically, we contrasted the diversity and compositional patterns obtained from metabarcoding and metagenomics, and assessed the concordance between their taxonomic and functional reconstructions. We further examined how different filtering strategies applied to metabarcoding data influenced the similarity between the two approaches, and critically evaluated the potential sources of observed biases. Finally, we tested how the different sequencing methods and filtering approaches affect the ecological interpretation and supported hypotheses. Based on these findings, we propose a set of guidelines to assist researchers in refining study designs and analytical strategies for investigating the gut microbiomes of bats and other similarly challenging taxa, such as birds.

## Methods

### Fieldwork

Fieldwork was carried out in spring-summer 2018 across the deserts of southern Israel (N 29.5-31.3, E 34.7-35.5), encompassing both the Negev Highlands and the Arava Valley. This area is characterised by low rainfall, between 30 mm and 220 mm (Evenari, 1981), and sparse vegetation cover, mostly restricted to wadis, temporary streams and rivers, which only run above ground during winter, and oases (Razgour et al., 2018). We captured bats with ground-level monofilament mist nets across this region. All species were identified using standard morphological measurements, and non-target species were released immediately at the place of capture. Individuals of the target species *Cnephaeus bottae*, *Hypsugo ariel*, and *Pipistrellus kuhlii* were retained in clean cloth bags for up to one hour and released at the capture place. Any faecal matter produced during this time was collected and stored immediately in absolute ethanol. All samples were frozen (-20°C) within 24 hours of collection and remained so until DNA extractions. The study was approved by the University of Southampton Ethics Committee, and was performed under permit 41282 of CK approved by the Israel Nature and National Parks Protection Authority.

### Laboratory sample processing

#### DNA extraction

DNA was extracted from one to three bat droppings (dry weight 10-20 mg) per bat using the PowerSoil® DNA Isolation Kit (MoBio, CA, USA). The manufacturer’s protocol was used with the modifications detailed in (Alberdi et al., 2018). Each extraction round included 23 bat faecal samples and one negative control. Subsequently, extracts were split for downstream processing through metabarcoding and metagenomics.

#### 16S rRNA amplicon sequencing metabarcoding

Metabarcoding was carried out on all sample extracts and all negative extraction controls. We used a broadly employed and validated primer set (341F, R806) (Caporaso et al., 2011; Muyzer et al., 1993) targeting the hypervariable V3 and V4 regions of the 16S rRNA gene to amplify the prokaryote barcode region. The addition of 24 different tags to the 5′-end of both primers enabled the differentiation of pooled samples after sequencing (Binladen et al., 2007).

All PCRs were set up in a dedicated pre-PCR laboratory to minimise contamination risk. A quantitative PCR (qPCR) screening on randomly selected extracts and blanks enabled us to assess contamination, determine optimal cycle numbers, and estimate the maximum DNA template volume that could be used without introducing PCR inhibition (Murray et al., 2015; Schnell et al., 2015). qPCR reactions were run on an Agilent Stratagene Mx3005P Thermocycler and an Applied Biosystems 2720 Thermal Cycler, using SYBR Green chemistry and a 25 µL reaction volume. Subsequently, each PCR reaction contained 2.5 µL AmpliTaq Gold buffer (1X), 2.5 µL MgCl□ (2.5 mM), 1.5 µL BSA (1.2 ng/µL), 0.5 µL dNTP mix (0.2 mM), 0.5 µL AmpliTaq Gold DNA polymerase (0.1 U/µL), 2 µL primer mix (0.8 mM), and 13.5 µL ddH□O. Thermal cycling conditions included an initial denaturation at 95°C for 10 minutes, followed by 30 cycles of 95°C for 15 seconds, 53°C for 20 seconds, and 72°C for 40 seconds, with a final extension at 72°C for 10 minutes. PCR product quality was verified via gel electrophoresis and GelRED staining under UV light. Amplicons were purified using an in-house SPRI bead protocol with 1:1 bead-to-DNA volume ratio to remove non-target DNA and primer dimers (DeAngelis et al., 1995; Rohland & Reich, 2012) and pooled based on band strength scores: 10 µL for low, 8 µL for medium, and 5 µL for high-intensity samples. Samples were grouped into batches of 24, ensuring each sample had a unique tag combination within the pool.

Sequencing libraries were constructed using the Blunt-End Single-Tube Tagsteady library building protocol (Carøe & Bohmann, 2020), with doubled adaptor concentrations. Libraries were quantified using an Agilent 2100 Bioanalyzer and sequenced on an Illumina MiSeq platform using paired-end 250 bp chemistry.

#### Shotgun metagenomics

DNA extracts were fragmented by ultrasonication on a Covaris LE220 platform, and sequencing libraries were prepared using the Blunt-End Single Tube (BEST) protocol with BEDC3 adaptors (Carøe et al., 2017). Library quality was assessed by qPCR on a Mx3005 qPCR System (Agilent, USA) using 1:20 diluted libraries. Each 25 µL qPCR reaction contained: 2.5 µL 10× PCR Gold Taq buffer, 2.5 µL MgCl□ (25 mM), 0.2 µL dNTP mix (10 mM each), 1 µL IS7 primer (10 µM), 1 µL IS8 primer (10 µM), 0.5 µL AmpliTaq GOLD DNA polymerase, 1 µL SYBR Green dye, 2 µL diluted library, and 14.3 µL sterile dH□O. Thermocycling conditions were: 95°C for 12 min; 40 cycles of 95°C for 20 s, 60°C for 30 s, and 72°C for 40 s; followed by a dissociation curve. Amplification curves were used to estimate the optimal number of PCR cycles for each library, minimising overamplification and clonality.

Sequencing libraries were then PCR-amplified with unique dual-index Illumina primers. Each 50 µL reaction included: 5 µL 10× PCR Gold Taq buffer, 5 µL MgCl□ (25 mM), 0.4 µL dNTP mix (10 mM each), 1 µL P7 primer (10 µM), 1 µL P5 primer (10 µM), 1 µL AmpliTaq GOLD DNA polymerase, 26.6 µL sterile dH□O, and 10 µL template library. PCR cycling conditions were: 95°C for 12 min; 7–20 cycles of 95°C for 20 s, 60°C for 30 s, and 72°C for 40 s; followed by 72°C for 5 min and a final hold at 4°C. The number of cycles per sample was determined from the preceding qPCR results. Indexed libraries were purified with SPRI beads at a 1:2 bead-to-library ratio, incubated for 5 min at room temperature, washed twice with 80% ethanol on a 96S Super Magnet, and eluted in EBT buffer at 37°C for 10 min. Final libraries were pooled and sequenced on an Illumina NovaSeq X platform, targeting 5 GB of data per sample.

### Bioinformatic analysis

#### 16S rRNA amplicon sequencing metabarcoding

Amplicon sequencing data were processed by demultiplexing reads according to molecular tag combinations using *AdapterRemoval* (Schubert et al., 2016), retaining only those sequences with exact matches to both tags and primers. Primer sequences were subsequently trimmed with *Cutadapt* (Martin, 2011). Quality filtering, trimming, and error modelling were carried out using *DADA2* (Callahan et al., 2016). Paired-end reads were merged with a minimum required overlap of 10 base pairs. Chimeric sequences were identified and removed prior to taxonomic classification, which was performed using the SILVA 138.1 reference database.

To reduce noise and potential false positives, the *Decontam* package (Davis et al., 2018) was used to identify and exclude likely contaminants. Additionally, ASVs classified as Eukaryota (ASV=1), Chloroplast (ASV=140), and Mitochondria (ASV=66) were removed from the dataset. Sample completeness was assessed using rarefaction curves generated with the ranacapa package (Kandlikar et al., 2018), and samples with fewer than 1,000 reads were excluded using *hilldiv2* (Alberdi & Gilbert, 2019b). ASVs representing less than 0.01% of total reads per sample were also filtered out.

#### Genome-resolved metagenomics

Raw sequencing reads were processed using the standard EHI bioinformatics pipeline (Eisenhofer & Alberdi, 2023). Initially, reads were quality-filtered with fastp (Chen et al., 2018), then aligned to the closest available reference genomes to our hosts using Bowtie2 (Langmead & Salzberg, 2012). The reference genome of *P. kuhlii* (GCF_014108245.1) was used for *P. kuhlii* and *H. ariel*, while the reference genome of *Eptesicus fuscus* (GCF_027574615.1) was used for *C. bottae*. Host-derived sequences were then removed from the resulting BAM files using samtools (H. Li et al., 2009). The remaining non-host reads were assembled both individually and collectively using MEGAHIT (D. Li et al., 2015).

To recover metagenome-assembled genomes (MAGs), binning was performed using a combination of CONCOCT (Alneberg et al., 2014), Maxbin2 (Wu et al., 2016), and MetaBAT2 (Kang et al., 2019), followed by refinement with the bin_refinement module from MetaWRAP (Uritskiy et al., 2018). MAGs were dereplicated at 98% average nucleotide identity (ANI) using dRep (Olm et al., 2017), leveraging MASH (Ondov et al., 2016) and FastANI (Jain et al., 2018) for similarity estimation. Non-host reads were then remapped to the dereplicated MAG catalogue using Bowtie2, and coverage statistics were calculated with CoverM (Aroney et al., 2025).

MAG quality was assessed using CheckM2 (Chklovski et al., 2022), and taxonomic classification was performed with GTDBtk v2.3.0 (Chaumeil et al., 2019) using the GTDB r214 database (Rinke et al., 2021). Functional annotation of MAGs was carried out with DRAM (Shaffer et al., 2020), integrating multiple reference databases (Bateman et al., 2004; Chan & Lowe, 2019; Finn et al., 2011; Huerta-Cepas et al., 2019; Kanehisa & Goto, 2000; Park et al., 2010; Seemann & Booth, 2018; Steinegger & Söding, 2017). Gene annotations were then translated into biologically interpretable Genome-Inferred Functional Traits (GIFTs) using distillR (Alberdi, n.d.). This tool uses a curated set of over 300 metabolic pathways and applies KEGG and Enzyme Commission (EC) identifiers to compute standardised GIFT scores ranging from 0 to 1. A score of 1 indicates the presence of all genes required for a pathway, 0.5 reflects partial presence, and 0 denotes complete absence.

#### Domain-adjusted mapping rate

When using genome-resolved metagenomics, the proportion of reads or bases mapped to the MAG catalogue is a common metric for assessing whether a microbiome has been adequately characterised (Marotz et al., 2018). However, when bacteria and archaea do not constitute the majority of DNA in a metagenome, or their representation varies considerably among samples, this metric can lead to misinterpretation by implicitly assuming that the prokaryotic fraction is larger than it actually is (Eisenhofer et al., 2024). The domain-adjusted mapping rate (DAMR) was developed to address this issue by normalising the mapping rate to the estimated bacterial and archaeal fraction rather than to the total metagenome (Pietroni et al., 2024).

To calculate DAMR, the prokaryotic fractions of metagenomes were estimated using SingleM’s Prokaryotic Read Fraction (SRF) function (Eisenhofer et al., 2024; Woodcroft et al., 2025) implemented in SingleM v0.16 with the S3.2.1.GTDB_r214_20231006 metapackage. Per-sample DAMR values were then calculated as:

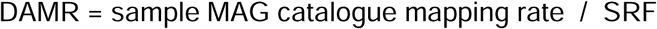

### Statistics

All statistical analyses were performed using R version 4.3.2 (R Core Team R, Others, 2013). Microbial alpha diversity was assessed using Hill numbers (Hill, 1973), which allow for the evaluation of different aspects of diversity—such as species richness, evenness, and phylogenetic relationships—across varying diversity orders. Specifically, we calculated species richness at q = 0 (presence/absence only), neutral diversity at q = 1 (weighted by relative abundance), and phylogenetic diversity at q = 1, using the *Hilldiv2* package (Alberdi & Gilbert, 2019b). To compare alpha diversity between species, we applied parametric t-tests when assumptions of normality and equal variance were met. In cases where these assumptions were violated, we used the non-parametric Wilcoxon test.

Beta diversity, or compositional dissimilarity between samples, was also calculated using Hill numbers. We computed Jaccard-type turnover for both neutral and phylogenetic beta diversity at q = 1 using the *hilldiv::hillpair function*. To visualise differences in microbial community structure, we generated nonmetric multidimensional scaling (NMDS) plots based on the resulting distance matrices. Dispersion differences between sampling methods were evaluated using the *betadisper* function from the *vegan* package (Oksanen et al., 2013). To test for overall compositional differences between groups, we conducted a PERMANOVA using *adonis2*, followed by pairwise comparisons with the *pairwiseAdonis* function (Martinez Arbizu, 2020).

Differential abundance of microbial taxa was analysed using ANCOM-BC2 (Lin & Peddada, 2024). To explore functional traits, MAGs were ordinated based on their Genome-Inferred Functional Traits (GIFTs) using t-SNE via the *Rtsne* package (Krijthe, 2015). Functional differences between communities were assessed by calculating community-weighted averages of GIFTs using *distillR::to.community*. Group comparisons were performed using the Kruskal-Wallis test, with Bonferroni–Holm correction applied to adjust for multiple testing.

#### Effects of data filtering

To better understand the differences between the metabarcoding and metagenomics data, we analysed the effects of data filtering in the recovery of ASVs. Specifically, we studied the individual and combined effects of two filtering criteria, namely copy number and prevalence. Copy number threshold overlooks the ASVs that recruit below a certain proportion of reads in each sample, while prevalence discards ASVs that are present in less than a given proportion of samples.

#### Functional characterisation

We analysed the impact of three data-generation strategies on proportion-type functional response attributes, measured across various traits, samples and species. The first employed standard processing of metabarcoding data (‘amplicon_standard’); the second also utilised metabarcoding but incorporated additional filtering to align as closely as possible with the metagenomic dataset (‘amplicon_filtered’); the third involved routine processing of genome-resolved metagenomic data (‘metagenomics’). We then fitted a beta-family generalised linear mixed model using the *glmmTMB* package (Brooks et al., 2017). Treatment was specified as a three-level fixed effect, while random intercepts for trait, sample and species accounted for nested and crossed grouping structure. Model convergence and goodness-of-fit were evaluated via residual diagnostics on the logit scale. Estimated marginal means and pairwise contrasts on the response (proportion) scale were computed using the *emmeans* package (Lenth et al., 2020), with Tukey adjustment for multiple comparisons.

## Results

### Data generation and DNA sequence fractions

Our metabarcoding data generation produced 4.5 million reads of the V3-V4 hypervariable regions of the 16S rRNA gene, yielding 87,222±62,466 amplicon reads per sample. These raw reads were processed into 3,852 amplicon sequence variants (ASVs). After initial filtering and decontamination, 3,250 ASVs were retained, spanning 27 distinct microbial phyla, including four archaea (Halobacterota, Thermoplasmatota, Euryarchaeota, Crenarchaeota) and 23 bacterial phyla (Fig. 1). The genome-resolved metagenomic approach was carried out based on 438.9 Gb (gigabases) of 150 bp paired-end sequencing data, ∼8.61 Gb per sample. The metagenomic assembly and binning yielded a catalogue of 135 dereplicated metagenome-assembled genomes (MAGs), spanning 13 bacterial and no archaeal phyla. The vast majority of ASVs (94.91%) and MAGs (72.59%) lacked species-level classification. Additionally, 32.13% of ASVs and 20.74% of MAGs lacked genus-level annotations.

**Figure 1.**
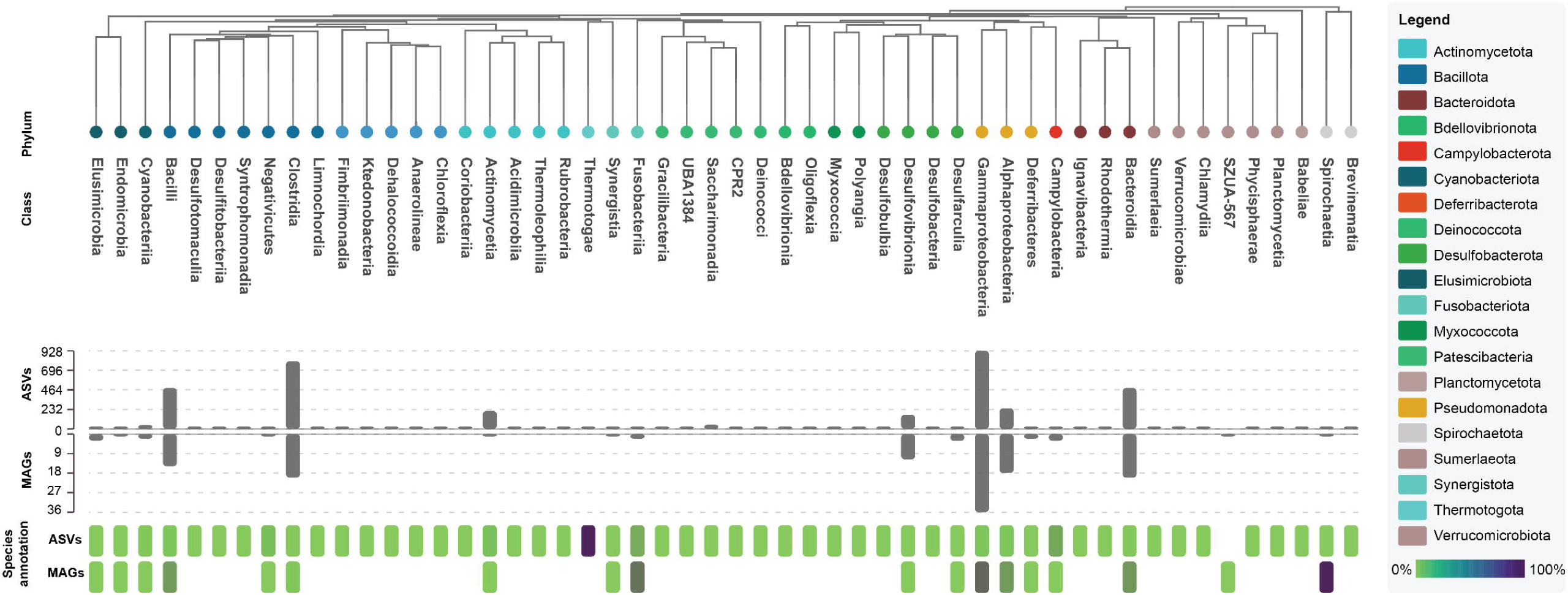
Phylogenetic tree of microbial classes captured through metabarcoding and metagenomics in the bat species *Pipistrellus kuhlii*, *Hypsugo ariel* and *Cnephaeus bottae* from the Negev Desert. Tip marker colors indicate the phyla listed in the legend box. Barplots indicate the number of amplicon sequence variants (ASVs) recovered through metabarcoding and the metagenome-assembled genomes (MAGs) reconstructed through metagenomics. Note the difference in scale. The species annotation heatmap shows the percentage of ASVs and MAGs with species-level annotation.

Unlike metabarcoding, metagenomics allows a more detailed examination of sequenced DNA fractions according to their host or microbiota origin, offering insights into the relative contribution of different data types in each sample. Our analysis revealed substantial variation in host-to-microbiota DNA fractions across species and samples (Fig. 2a). The average estimated microbial fraction was 22.1±19.6%, although this varied significantly (K-W: X^2^ = 18.395, p = 0.0001) across species, with *H. ariel* carrying lower microbial fractions than the other species. Substantial variation was also observed across samples within species, with host-mapped reads ranging from 1 to 75% and reads mapped against the MAG catalogue spanning from 1 to 86%. We estimated the success of microbial recovery by normalising MAG catalogue-mapped reads with the total estimated microbial reads, yielding the domain-adjusted mapping rate (DAMR). Across all samples, the dataset showed an average DAMR of 84.9 ± 20.6, with the highest values in *P. kuhlii* (90.3 ± 17.1), followed by *H. ariel* (83.8 ± 18.6) and *C. bottae* (77.9 ± 28.2) (Kruskal–Wallis: p < 0.01). In most cases, the mapped fraction was comparable to or exceeded the estimated microbial fraction—indicative of good microbial recovery, although seven samples showed notable discrepancies between estimated and mapped values (Fig. 2b). While the estimated prokaryotic fraction was positively associated with metagenomics-derived neutral alpha diversity (Spearman’s: S = 10011, p = 0.001), we observed no significant relationship with diversity metrics of metabarcoding data (Spearman’s: S = 16198, p = 0.99) (Fig. 2c).

**Figure 2.**
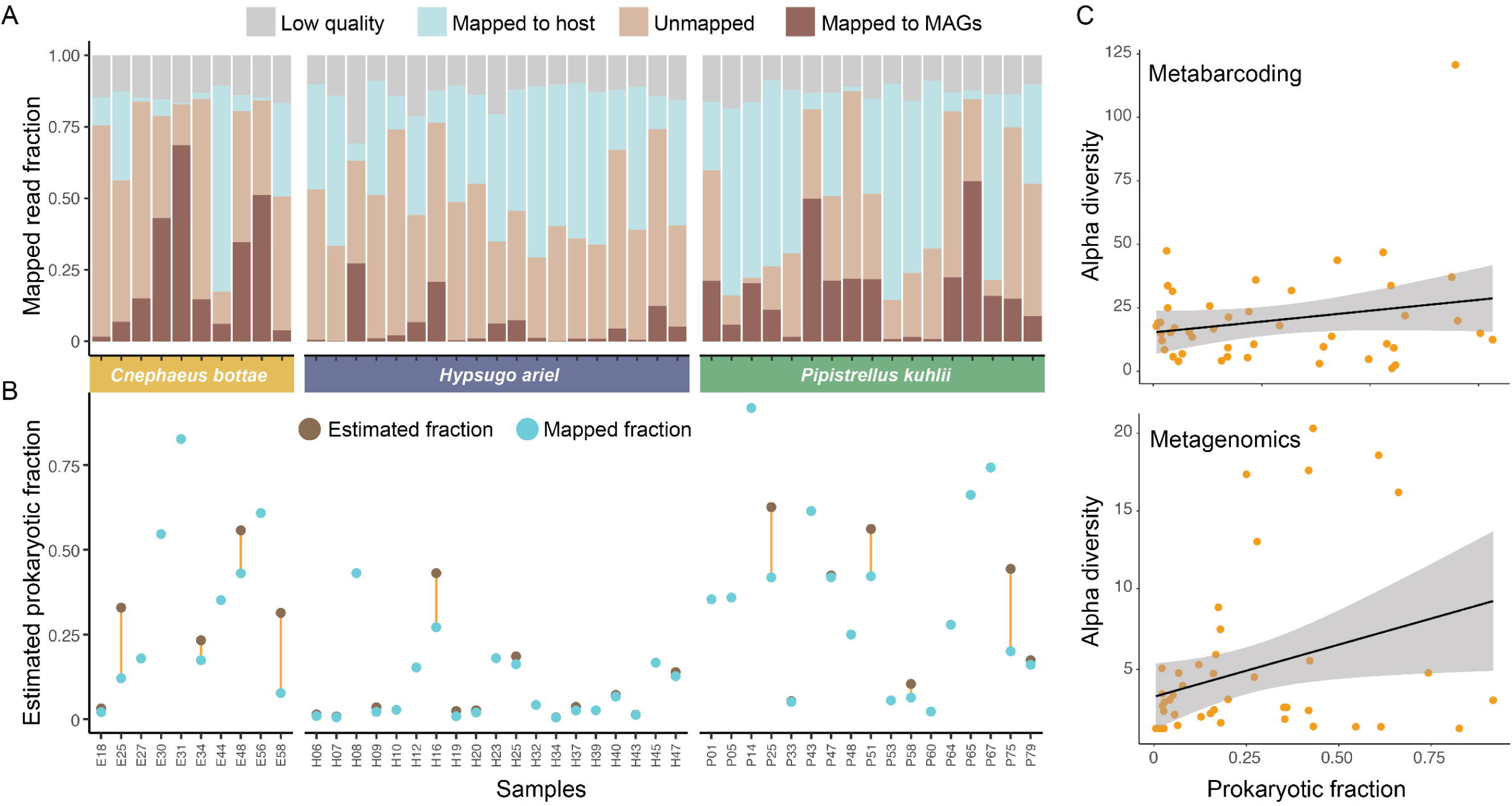
DNA fractions and estimated microbial recovery in the metagenomics dataset. **A)** Proportions of metagenomic reads discarded due to low quality, mapped to the host genome, mapped to the MAG catalogue and unmapped to any reference. **B)** Difference between the prokaryotic fraction estimated based on marker genes and mapped fraction to the MAG catalogue. The orange line represents the gap between both metrics, indicating the estimated prokaryotic fraction missed in each sample. **C)** Relationship between estimated prokaryotic fraction and the measured alpha diversity in terms of neutral diversity (Hill number of q=1 or exponential of Shannon index) for 16S amplicon sequencing (metabarcoding) and genome-resolved metagenomics (metagenomics) datasets.

### Taxonomical characterisation

Each bat species exhibited a distinct microbial community composition. Across all samples, most ASVs (3387, 88.34% of the total) and MAGs (82, 60.74%) were unique to a single host species. Among these, *Cnephaeus bottae* harboured the largest proportion of species-specific ASVs (79.43%) and MAGs (53.26%). In terms of abundance, *Pipistrellus kuhlii* contained the highest number of ASVs (1523), followed closely by *Hypsugo ariel* (1500) and *C. bottae* (1381). Conversely, the genome-resolved metagenomics dataset revealed the highest number of MAGs in *C. bottae* (92), followed by *P. kuhlii* (69) and *H. ariel* (30). Additionally, a large fraction of ASVs were individual-specific. Overall, 77.70% of ASVs (2979) were detected in only a single individual, with *H. ariel* exhibiting the highest proportion of unique individual ASVs (70%). In contrast, only 25.19% of MAGs were found in a single individual, although *H. ariel* again showed the highest proportion (30%)(Supplementary Table 2).

At the phylum level, differences among bat species were also evident. Amplicon sequencing revealed that *P. kuhlii* and *H. ariel* samples contained the highest numbers of bacterial (20 and 19, respectively) and archaeal phyla (4 and 3), whereas *C. bottae* showed the lowest (17 bacterial and an archaeal phylum; Supplementary Table 1-2). Fifth phyla (Actinomycetota, Cyanobacteriota, Planctomycetota, Patescibacteria, Synergistota) showed significant differences in their abundances across species (Supplementary Table 7a). In contrast, metagenomic data indicated that *C. bottae* hosted the highest number of bacterial phyla (10), while *H. ariel* exhibited the lowest (5 phyla; Supplementary Table 1 & 3). Differential abundance analysis only found differences in Bacteroidota, where *H. ariel* showed significantly lower abundance than the other two species (p adjusted<0,05; Supplementary Table 7b). However, both methodological approaches identified Pseudomonadota as the most abundant phyla in the gut microbiota of the bats, followed by Bacillota and Bacteroidota. The main differences were in their capacity to detect less abundant bacteria (Supplementary Table 1).

Whereas no archaeal MAG was reconstructed through genome-resolved metagenomics, a total of 39 archaeal ASVs were recovered through metabarcoding. These ASVs belonged to four phyla and were identified in 16 individuals across the three species, with an average of 1.1±3.2 ASVs per individual. The majority of archaeal ASVs were found in *H. ariel* (24 ASVs), followed by *P. kuhlii* (19 ASVs) and *C. bottae* (2 ASVs). However, most were detected in only a single sample (87%), with only four individuals harbouring more than one archaeal ASV. A single *H. ariel* sample stood out, containing 20 archaeal ASVs. Although archaeal ASVs were found across all three species, they were more prevalent in *P. kuhlii* (7 ASVs) and *H. ariel* (5) than in *C. bottae* (2). The 39 ASVs were classified into 11 genera, with all genera present in more than one individual. Among them, the most frequent ones were *Methanobrevibacter* from phylum Euryarchaeota, and *Halostagnicola* (5), and *Haloterrigena* (4), Methanimicroccus (4) from Halobacterota phylum.

### Microbiome diversity and composition

Due to the impossibility to match most ASVs and MAGs at species and genus level, we calculated alpha diversities at two higher taxonomic resolutions: family and phylum levels (see supplementary Tables 5). At the family level, the methodological approach explained all the the variabiliy in richness (53%, F(1) = 112.37, p < 0.0001) and in neutral diversity (27%, F(1) = 35.42, p < 0.0001). At the phylum level, both *species* (F(2) = 5.20, p = 0.007; F(2) = 7.39, p = 0.001) and *method* (F(1) = 94.92, p < 0.001; F(1) = 14.85, p < 0.001) had significant effects on both alpha diversity metrics. In both cases, the methodological approach accounted for a greater proportion of the variance than host species. No significant interaction between *species* and *method* was detected for alpha diversity metrics at either taxonomic level. When analysing the pattern observed with each methodological approach, *C. bottae* consistently exhibited the highest average diversity values in both metagenomic and metabarcoding analyses. However, metabarcoding did not detect differences between species for any diversity metrics at either taxonomic level, whereas metagenomics did. At family level, *C. bottae* showed significantly higher richness than the other two species (*H. ariel*, p=0.0008, *P. kuhlii* p=0.049), while differences in neutral diversity were only observed between *C. bottae* and *H. ariel* (p = 0.0316), with H. ariel displaying markedly lower diversity in the metagenomic dataset. At the phylum level, these differences became more pronounced: richness varied significantly among the three species (p < 0.001), and the neutral diversity metric revealed lower diversity in *H. ariel* compared to both *C. bottae* (p = 0.004) and *P. kuhlii* (p = 0.005).

Similarly, we compared the beta diversity at family and phylum level. At the family level, a PERMANOVA using methodological approach and species as explanatory variables revealed a significant overall effect of the model on community composition, the effect being higher in richness (F = 4.85, R² = 0.20, p = 0.001) than in neutral metric (F = 2.98, R² = 0.13, p = 0.001). The explanatory power of the methodological approach decreased from richness (12%) to neutral metric (5%; Supplementary Table 6). At the phylum level, the variability explained by the methodological approach increased to 22% in the richness (F = 5.00, R² = 0.22, p = 0.001), while in neutral metric only the *species* explained the variability (F = 5.66, R² = 0.10, p = 0.003; Supplementary Table 6). Their interaction was significant only in richness at both taxonomic levels (F = 2.04, R² = 0.03, p = 0.001; Phylum, F = 5.62, R² = 0.07, p = 0.001). The permutation test for homogeneity of multivariate dispersions indicated a significant difference in dispersion among groups at family level (F = 7.70, p = 0.002), suggesting some variation in within-group community variability across species and methods. We further analysed these discrepancies between the methodological approaches by analysing the compositional changes within each method. At family level, richness-based metrics detected differences only between *H. ariel* and the other two species in both methodological approaches. When applying the neutral beta diversity metric, in metabarcoding, only *C. bottae* was the one that differed from *H. ariel* (p = 0.003), whereas in the metagenomics approach the differences were found between *P. kuhlii* and *H. ariel* differed significantly (p = 0.015). At phylum level, the patterns diverged, with metagenomics identifying a greater number of interspecific differences than metabarcoding (Supplementary Table 8).

### Effects of data filtering

To gain insights into the source of the mismatches observed between metabarcoding and metagenomics datasets, we applied multiple filtering criteria to the raw metabarcoding dataset (3852 ASVs) based on minimum prevalence and fraction of reads. The number of ASVs decreased drastically when applying both filtering thresholds, dropping to 79 when applying the most stringent read fraction threshold, and 53 when applying the most stringent prevalence threshold. Although 50 ASVs were identified as archaea, applying even a mild prevalence threshold of 10% reduced this number to just three. A read fraction threshold of 0.01 was required to produce a similar reduction. Diversity metrics were also significantly affected by these thresholds. As expected, the diversity values decreased drastically in the three species when applying both filters. Total richness dropped from 118.84 ± 111.46 to 7.88 ± 7.41, and neutral diversity declined from 20.65 ± 20.62 to 3.06 ± 2.17 under the most stringent copy number threshold. Applying the strictest prevalence threshold further reduced these values to 2.37 ± 0.92 and 2.24 ± 0.89, respectively (Fig. 4a).

Dissimilarities in diversity between the two approaches disappeared at a prevalence of 40% without any read fraction cut-off. When applying cut-off thresholds, a minimum copy number threshold of 0.01 and a prevalence of 10% were necessary for convergence. The metabarcoding filtering thresholds that best matched metagenomic alpha diversity were a cut-off of 0.01 with a prevalence of 10% (distance = 0.77) or no read fraction cut-off with a prevalence of 30% (distance = 1.10). The NMDS plots show that the filtering threshold cut-off 0.01 and a prevalence threshold of 10% had higher convergence with the metagenomics dataset (Fig. 4b). We then compared the differences in community composition between the filtered metabarcoding and the metagenomics datasets to understand the influence of rare ASVs in the results. With a cut-off of 0.01 and a prevalence threshold of 10%, metabarcoding still yielded more types (165 ASVs) than metagenomics (135 MAGs) although spanning only 9 phyla, compared to the 13 phyla recovered in metagenomics (Supplementary Table 4).

After filtering, the species ranking by ASV count changed: *C. bottae* became the species with the highest number of ASVs (130), similar to the pattern observed in metagenomics, followed by *H. ariel* (45) and *P. kuhlii* (19). Despite both approaches detecting the same dominant phyla (Fig. 3A; Supplementary Table 1), metabarcoding showed a lower relative abundance of Fusobacteriota. These differences were primarily driven by *C. bottae* and *P. kuhlii*, where metagenomics detected more Fusobacteriota. Moreover, metabarcoding detected more Bacillota in *H. ariel*.

**Figure 3.**
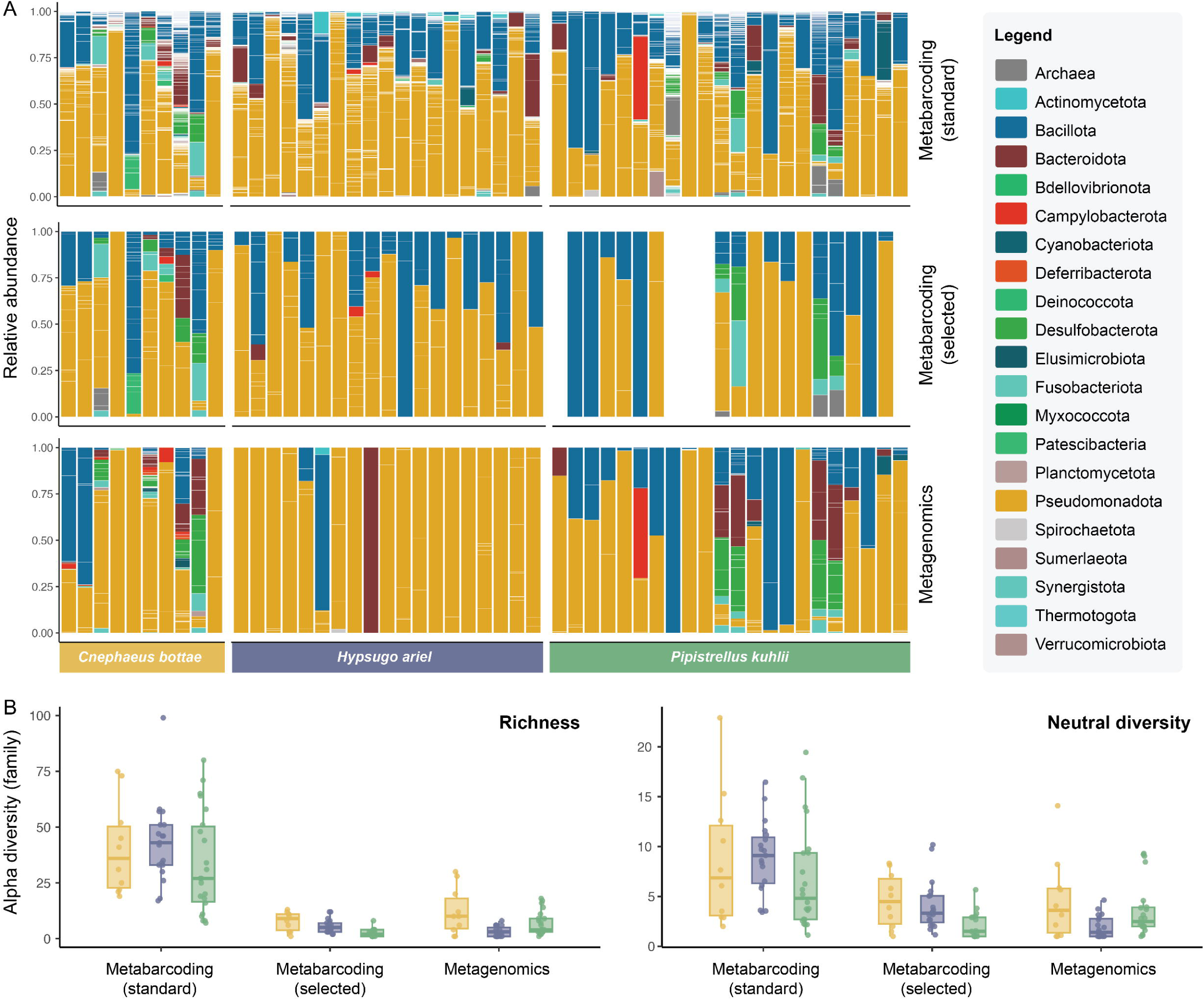
Microbiome composition and diversity recovered by the different data generation and processing approaches. **A)** Taxonomic composition of the microbiome recovered through metabarcoding with standard filtering, metabarcoding with selected filtering and metagenomics. The empty bars in Metabarcoding (selected) are due to complete removal of data in certain samples following the filtering criteria. **B)** Family-level alpha diversity based on richness (presence-absence of ASVs or MAGs) and neutral diversity (Hill number of q=1 or exponential of Shannon index) metrics for the three data generation and processing approaches.

Filtering rare taxa made alpha diversity estimates more comparable between methods (Fig. 3B; Supplementary Table 5). Post-filtering, species identity significantly explained variation in both richness and neutral diversity metrics (p < 0.001), whereas the methodological approach alone did not (p > 0.1). However, a significant interaction between species and method (p < 0.05) indicated that methodological effects varied among host species. At the family level, both diversity metrics revealed significant differences between *P. kuhlii* and the other two species (p < 0.05), whereas at the phylum level, richness differed between *C. bottae* and the remaining species (p < 0.05), but no significant differences were detected in the neutral metric.

Regarding beta diversity, both species and method significantly explained variability in community composition at both taxonomic levels and for both diversity metrics (Fig. 4; Supplementary Table 6). Species identity accounted for a greater proportion of variance than methodological approach, with slightly higher explanatory power observed at the phylum level (richness: 16%, neutral: 10%) compared to the family level (both metrics: 7%).

**Figure 4.**
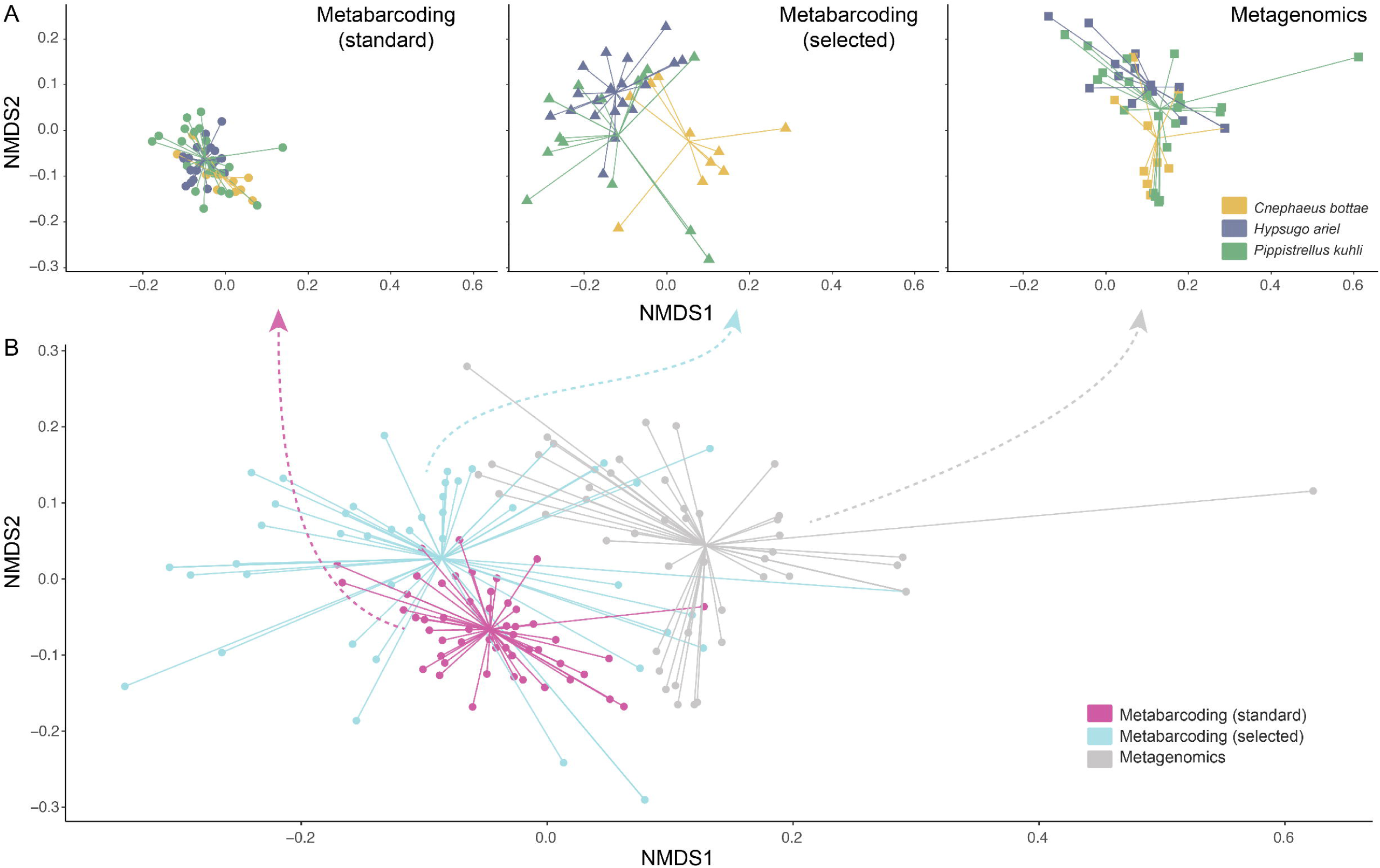
Compositional differences between the three bat species and methodological approaches. **A)** Compositional variation faceted by methodological approach, with samples coloured by host species. **B)** Compositional variation across samples for the entire dataset, with samples coloured by methodological approach. Dissimilarities are computed using pairwise beta diversities at family level.

### Functional insights from different datasets

Lastly, we characterised the functional attributes of bacteria: directly through gene annotation of MAGs in the case of metagenomics, and indirectly through reference-based functional inference in metabarcoding. The characterisation of functional attributes of the microbiome yielded significantly different results based on the type of dataset and filtering strategy employed (Fig. 5). The back□transformed estimated marginal means of the functional traits were highest for the amplicon□filtered treatment (0.481; SE = 0.0306; 95% CI = 0.422–0.541), followed closely by the amplicon□standard approach (0.469; SE = 0.0306; 95% CI = 0.410–0.529), and lowest for metagenomics (0.346; SE = 0.0277; 95% CI = 0.294–0.402). Both amplicon□based workflows produced significantly higher estimates of metabolic capacities than metagenomics (OR = 1.75 and 1.67; p < 0.0001), while the difference between the two amplicon methods was statistically significant but modest (OR = 1.05, p = 0.043) (Fig. 5).

**Figure 5.**
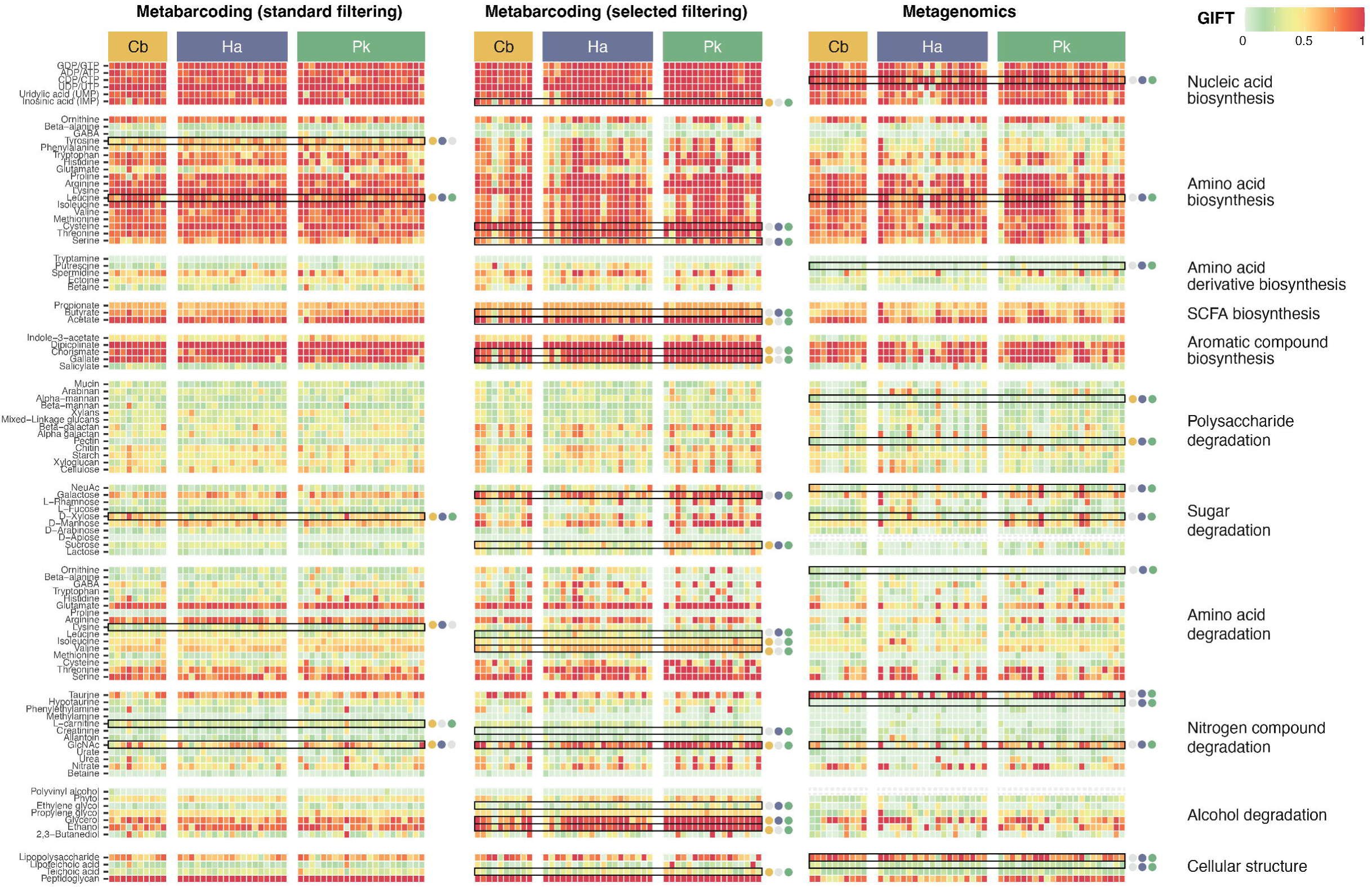
Genome-inferred functional trait (GIFT) differences across species and methodological approaches. Black boxes indicate statistically significant differences between species, with the coloured dots indicating the species between which significant differences were measured.

Across methods, we detected substantial variation in the number and identity of functional traits differing among bats. The metabarcoding standard revealed differences in only eight functions, with a comparable range of differences across species pairs (2–4 functions). In contrast, metagenomic profiling identified a total of 19 traits that differed among species, with the greatest separation observed between *H. ariel* and *P. kuhlii* (16 functions), and the smallest between *P. kuhlii* and *C. bottae* (one function). The metabarcoding-selected approach detected the highest number of functional differences overall. Similar to the metagenomic results, the largest functional divergence involved 16 traits, however, in this case the greatest differences were found between *P. kuhlii* and *C. bottae*, rather than between *P. kuhlii* and *H. ariel*. Even so, the latter species pair still differed in 10 functions. In general, the lowest variability across methodological approaches was consistently observed between *C. bottae* and *H. ariel*.

Beyond differences in the number of traits identified, the specific functions contributing to these differences varied markedly among species and methods. This high functional turnover prevented the identification of consistent functional patterns shared across species comparisons.

## Discussion

Animal-associated microbiomes can be investigated using various approaches, each defined by specific technical steps and limitations that can substantially influence the results and their interpretation (Aizpurua et al., 2023; Zepeda Mendoza et al., 2015). In this study, we present a comparative analysis of two widely used microbiome profiling techniques, metabarcoding (16S rRNA amplicon sequencing) and genome-resolved shotgun metagenomics, to characterise the gut microbiota of three bat species. Our comparison revealed marked differences in the recovered microbial communities, as well as in their taxonomic and functional profiles, and provided insights into potential sources of methodological bias. We found that these differences shaped the ecological interpretation of the study and the supported hypotheses, underscoring the importance of critically evaluating microbiome data in light of methodological constraints. Overall, this work offers a comprehensive assessment of the strengths and limitations of metabarcoding and metagenomic approaches for profiling gut microbiomes, with implications for the design and interpretation of studies in bats and other taxa with low-biomass, high-variability samples, such as birds.

### Compositional microbiome recovery through metabarcoding and metagenomics

Metabarcoding and metagenomics yielded similar coarse-level microbiota reconstructions, with strong congruence in terms of the most abundant bacterial phyla found in the gut microbiota of the three bat species and the bacterial phyla with >1% relative abundance. Both methods identified Pseudomonadota (=Proteobacteria) as the most abundant phylum (overall average relative read abundance 59% and 68%, respectively), followed by Bacillota (=Firmicutes) (26% and 16%) and Bacteroidota (5% and 7%). These results are in line with previous bat microbiome characterisations conducted across multiple taxa in China (Dai et al., 2024), India (Selvin et al., 2019), North America (Arellano-Hernández et al., 2024), and Europe (Aizpurua et al., 2021; Vengust et al., 2018).

The differences between methods became increasingly pronounced with deeper taxonomic resolution, showing almost no congruence at the family level. While discrepancies between the reference databases used for annotating metabarcoding and metagenomic data likely explain part of this variation (Wright et al., 2023), given frequent taxonomic revisions and nomenclatural inconsistencies following phylogenetic reclassification (Oren & Garrity, 2021; Parks et al., 2018, 2020), the extent of diversity captured by each method also played a major role. Unfiltered amplicon data suggested a microbial community 26.2 times more diverse than that recovered by metagenomics, including 50 ASVs spanning four archaeal phyla that were entirely absent from the metagenomic dataset. These discrepancies likely reflect a combination of diversity overestimation by metabarcoding combined with underestimation by genome-resolved metagenomics.

Metabarcoding can surface the signal of low-abundance taxa better than genome-resolved metagenomics through the targeted amplification and sequencing of the 16S rRNA gene in bacteria and archaea. However, unless explicit corrections are applied, this method considers each sequence variant of the 16S rRNA gene as a different type in the diversity analyses (Alberdi & Gilbert, 2019a), often interpreted as a different strain (Johnson et al., 2019). As prokaryotic genomes usually contain multiple copies of the 16S rRNA gene, often with slightly different sequences, this approach can lead to an inflation of diversity (Větrovský & Baldrian, 2013). In addition, PCR amplification and sequencing errors can introduce artefacts that, despite the algorithmic efforts to discard them (Callahan et al., 2016; Davis et al., 2018), can further inflate diversity estimation (Nearing et al., 2021; Roume et al., 2023). Supporting this interpretation, we found that over 70% of ASVs identified through amplicon sequencing were exclusive to single individuals, whereas only 25% of the genomes were exclusive to a single individual in the metagenomic dataset.

In contrast, genome-resolved metagenomics tends to underestimate diversity due to the technical constraints inherent in genome reconstruction (Malmstrom & Eloe-Fadrosh, 2019). This approach depends on the presence of sufficient bacterial reads, ideally distributed across multiple samples, to enable the algorithmic recovery of genomes through differential coverage of assembled contigs (Albertsen et al., 2013). As a result, low-abundance and rare taxa that fail to generate enough coverage for assembly are often overlooked (Eloe-Fadrosh et al., 2016). Moreover, the stringent preprocessing criteria typically applied in genome-resolved metagenomics to minimise false positives, such as excluding metagenome-assembled genomes (MAGs) with completeness below 50% (Bowers et al., 2017), or requiring a minimum genome coverage of 30% for valid detection (Zhao et al., 2023), can further limit the recovery of the full microbial community. These constraints likely contributed to the lower detection rates of archaeal taxa and individual-specific variants observed in our study relative to amplicon sequencing.

Our results indicate that the biases and limitations of genome-resolved metagenomics are particularly pronounced in challenging sample types such as bat guano. DAMR analyses showed that, in most cases, the mapping rate to the reference genome catalogue closely matched the microbial fraction estimated from single-copy core genes. Furthermore, the finding that doubling the sequencing depth produced only marginal gains in recovered microbial diversity suggests that sequencing effort was not the primary limiting factor for microbiome characterisation. Instead, the combination of low average microbial DNA content and high variability among samples likely constrained the successful reconstruction of bacterial genomes and, consequently, downstream microbiome profiling. Genome-resolved metagenomics failed to recover most low-abundance taxa detected by metabarcoding, an effect most pronounced in *H. ariel*, the species with the lowest microbial DNA fractions. The fact that the greatest congruence between methods was achieved when the metabarcoding dataset was filtered to include only ASVs with a minimum relative abundance of 1% and a minimum prevalence of 10% of samples further supports this interpretation. Previous studies showed that adding host DNA to mock bacterial communities decreases metagenomic sequencing coverage for characterising the microbiome, with sensitivity to detect low abundant bacterial taxa decreasing with increasing proportion of host DNA (Pereira-Marques et al., 2019).

### Functional profiling through metabarcoding and metagenomics

The limitations of genome-resolved metagenomics in low-microbial-biomass samples were further highlighted by the functional analyses. Although this approach is often praised for its ability to provide direct functional insights through the annotation of reconstructed genomes, individual functional inferences are strongly influenced by genome completeness (Eisenhofer et al., 2023), and undetected taxa can further distort community-level functional profiles. In our study, the average metabolic capacity of microbial communities was consistently lower when assessed directly from genome-resolved metagenomics than when inferred indirectly from metabarcoding data. This discrepancy likely reflects that functional inference tools such as PICRUSt rely on complete or near-complete reference genomes (Douglas et al., 2020), whereas our reconstructions included many partially assembled genomes.

A major criticism of indirect functional inference, particularly in studies of wild taxa, is that reference genomes often originate from bacteria isolated from humans or other model organisms, and thus may not accurately represent the functional repertoire of environmental bacterial communities (Djemiel et al., 2022; Matchado et al., 2024; Sun et al., 2020). Because both our metabarcoding-based functional inferences and metagenomic taxonomic annotations drew from the NCBI reference genome database, we were able to directly match reference genomes used for ASV-based functional inference with 30 of our 140 reconstructed genomes. The comparison of their gene annotations revealed broadly similar functional patterns, with expected discrepancies in low-completeness genomes. A few reconstructed genomes exhibited unique metabolic traits absent from reference genomes, including pathways involved in amino acid derivative biosynthesis, amino acid and nitrogen compound degradation, and xenobiotic and antibiotic metabolism, but differences were minor.

However, this collection of 30 matches represented only 21.4% of the reconstructed genomes, the subset showing the highest similarity (<3% average nucleotide dissimilarity) to reference genomes. Consequently, they correspond to the fraction expected to be functionally most comparable to existing references. The remaining reconstructed genomes showed average nucleotide identities below 95% when compared to GTDB representative genomes, the commonly accepted threshold for microbial species delineation (Jain et al., 2018). Extrapolating these proportions to ASVs suggests that nearly 80% of ASVs were assigned functional traits based on distantly related reference genomes, likely reducing the reliability of functional predictions, potentially by a substantial degree.

### Effect of result discrepancies on ecological inferences

Bats represent approximately 20% of mammalian diversity, with over 1,500 species currently described (Simmons & Cirranello, 2020). Owing to their key roles in ecosystem functioning, bats have been proposed as valuable indicators of biological, ecological, and environmental disturbance (Jones et al., n.d.). Furthermore, their contribution to essential ecosystem services, such as pest control (Kunz et al., 2011), makes them a compelling system for exploring how gut microbiomes may influence ecological processes.

The methodological limitations and biases were reflected in different alpha and beta diversity metrics, yielding conflicting ecological conclusions depending on the employed methodology. For instance, metabarcoding data suggested *H. ariel* possessed a distinct microbiota, thus supporting the shared habitat hypothesis, whereas shotgun metagenomics identified *P. kuhlii* as the most distinct taxon, supporting the shared evolutionary history hypothesis. It is interesting that contrary to our shared diet hypothesis, neither approach identified *C. bottae* as having a distinct gut microbiota despite its unique diet. While the diet of both *P. kuhlii* and *H. ariel* mainly comprises Diptera and Lepidoptera, *C. bottae* primarily feeds on Hymenoptera and Lepidoptera (Feldman et al., 2000). In contrast, previous studies of bat microbiomes found correlations between the gut microbiome and diet composition of eight insectivorous bats in China (Dai et al., 2024).

*C. bottae* was identified as having the lowest number of bacterial phyla and ASVs by the metabarcoding approach, but the highest number of bacterial phyla and MAGs based on metagenomics. Despite these quantitative differences, the overall community structure at the level of dominant phyla remained relatively consistent between *C. bottae* and *P. kuhlii* across both methodological approaches, with discrepancies primarily confined to low-abundance taxa (<0.2%). However, *H. ariel* represented a notable exception, showing pronounced differences in the abundance of two major phyla: Bacteroidota and Pseudomonadota, thus supporting our shared habitat hypothesis. While metagenomic data highlighted Bacteroidota as the principal phylum distinguishing *H. ariel* from the other two species, metabarcoding did not capture this difference. Although we cannot determine whether this difference is real, it highlights how methodological choices can lead to contrasting interpretations. This is particularly important because Bacteroidota are recognised as key gut bacteria involved in the degradation of complex polysaccharides and gut physiology (Thomas et al., 2011). Thus, methodological biases that obscure or exaggerate differences in Bacteroidota abundance may lead to misleading ecological inferences regarding host dietary preferences (the ratio of Lepidoptera/Diptera) or gut transit in *H. ariel*.

After applying more stringent filtering criteria (a cut-off of 0.01 and a prevalence threshold of 10%), the total number of ASVs was drastically reduced, as was the number of ASVs detected in each species, largely due to the high proportion of ASVs occurring in only a single individual. Interestingly, *C. bottae* shifted from being the species with the lowest number of ASVs to the highest, followed by *H. ariel* and *P. kuhlii*. The number of phyla and families identified in each species also followed this same ranking pattern observed for ASV counts. Consequently, alpha diversity values decreased after filtering. Under standard filtering, no significant differences in alpha diversity were observed among species; however, after applying the more stringent filtering, *C. bottae* exhibited higher diversity than the other two species. In terms of beta diversity, the stricter filtering increased the proportion of variance explained by species identity (11% compared to 6%), and pairwise comparisons revealed significant differences among all species, whereas no such differences were detected under the standard filtering. These results demonstrate that not only the methodological approach used for data generation but also the analytical choices made during data processing can profoundly influence the outcomes. This underscores the importance of thorough data evaluation and sensitivity testing to ensure robust and biologically meaningful interpretations.

The functional analysis produced contrasting patterns depending on the sequencing method and filtering approach. Contrary to previous functional studies warning that metabarcoding approaches can overestimate functional divergence (Louca et al., 2018), the standard metabarcoding approach in our study yielded the lowest number of functional differences among the gut microbiomes of the three bat species with no pattern of stronger differentiation between any species pairs. However, the filtered (selected) metabarcoding approach yielded the highest number of differentiated functions, highlighting the impact of filtering on ecological inference. The filtered metabarcoding approach identified the highest number of functional differences between *P. kuhlii* and *C. bottae* and lowest between *C. bottae* and *H. ariel,* thus supporting our shared evolutionary history hypothesis. In contrast, the metagenomics approach supported the shared habitat hypothesis, with greatest functional differences identified between the microbiomes of *H. ariel* and *P. kuhlii*, followed by *H. ariel* and *C. bottae*. Previous studies assigned functional shifts in the gut microbiome to changes in diet (Gibson et al., 2019). In our study, changes in diet can explain the shared habitat hypothesis as different prey items may be available in anthropogenic and natural habitats.

### Practical considerations for microbiome analysis of low-biomass, highly variable gut microbiomes

The biological discrepancies between the results yielded by the different methods highlight the importance of critically assessing the results. Different study systems and datasets might call for different methodologies, but this requires acknowledging the biases and limitations of the chosen methodologies.

Genome-resolved metagenomics is often regarded as superior to 16S rRNA amplicon sequencing for microbiome profiling because it enables direct functional inference from reconstructed genomes, while overcoming taxonomic biases of PCR amplification of marker genes. However, most studies supporting this view are based on hosts such as humans, laboratory mice, or livestock, which produce faecal samples enriched in bacterial and archaeal DNA with relatively stable microbial biomass. A recent survey of over 150 wild vertebrates revealed that non-flying mammals, reptiles, and amphibians typically harbour a comparable microbial fraction of around 75%, with slightly lower values in strict carnivores and higher ones in omnivores and herbivores due to dietary digestibility ((Pietroni et al., 2024)). When more than three-quarters of sequencing reads derive from microbial DNA, the probability of capturing sufficient coverage for genome reconstruction is high, often resulting in hundreds of high-quality MAGs even with moderate sequencing effort. For some comparative insights, three times less sequencing than in our study yielded 539 MAGs from Pyrenean brook salamander (*Calotriton asper*) faecal samples, and with half of the sequencing effort of this study, 860 MAGs were produced in the European hare (*Lepus europaeus*) (Aizpurua et al., 2024). In contrast, our analysis of bat guano produced only 140 MAGs despite much deeper sequencing, with microbial DNA fractions averaging around 22.1%, similar to those reported for bats and birds in (Aizpurua et al., 2024)).

This reality raises legitimate questions about the suitability of genome-resolved metagenomics for gut microbiome studies in such taxa. When microbial DNA represents only 5% of total reads, the remaining 95%, mostly host and dietary DNA, must also be sequenced to achieve adequate microbial coverage, substantially increasing costs. Researchers should therefore carefully assess whether genome-resolved metagenomics is the most appropriate approach for their study goals or whether metabarcoding could provide sufficient insight. If a primarily taxonomic characterisation is sufficient, metabarcoding may offer a more practical and realistic representation of microbial communities, provided that stringent filtering criteria are applied to minimise false positives.

While low microbial fractions can, in principle, be offset by increasing sequencing depth, the second major challenge—high variability in microbial load among samples—is harder to address. Our results yielded a few samples of *P. kuhlii* and *C. bottae* exceeding 75% of estimated microbial fraction, which align with previously reported observations of microbial fraction spikes above 95% in a few bat and bird individuals (add ref.). One must therefore consider whether a microbiome profile derived from an individual with 95% microbial DNA is comparable to that from an individual with only 5%. Recent work indicates that host DNA shedding tends to remain stable (Tang et al., 2025), suggesting that fluctuations in microbial fractions primarily reflect variable microbial biomass. Incorporating estimated microbial biomass as a covariate in statistical analyses, similar to how sequencing depth is accounted for, may help mitigate this bias. However, in hosts with extremely rapid digestion, such as bats, whose intestinal passage time is typically between 25 and 204 minutes, depending on body mass and diet (Cornelius Ruhs et al., 2024; Roswag et al., 2012), dietary DNA can represent a substantial proportion of total faecal DNA, complicating such corrections.

Although 16S amplicon sequencing might appear less sensitive to these biases, many of the same limitations still apply, albeit invisibly to researchers. Our results show that metabarcoding can indeed better capture signals from low-abundance and low-prevalence taxa than genome-resolved metagenomics, yet the variation in microbial DNA fractions across samples persists. Consequently, the number of microbial template molecules available for PCR amplification varies substantially, likely influencing diversity estimates. Hence, for studies dealing with low-biomass and highly variable samples, quantitative PCR (qPCR) assays with reference standards are strongly recommended to estimate initial bacterial loads and incorporate this information into downstream analyses.

Functional inference presents an additional layer of complexity. Indirect functional predictions from 16S data in wild animals are often criticised because they rely on reference genomes characterised from different environments, which may carry distinct functional repertoires (Albright & Louca, 2023; Matchado et al., 2024). Conversely, genome-resolved metagenomics may underestimate functional diversity when many reconstructed genomes are incomplete. When species-level annotation rates are high, reference genomes of closely related taxa may provide reliable functional inference. However, in systems where most ASVs lack species-level assignment, as is common in wild microbiomes, functional predictions are likely to be distorted. Indirect functional inference may suffice for broad comparisons, such as functional diversity metrics, but is less appropriate for detailed pathway-level analyses. Thus, researchers should evaluate which methodological limitations are most likely to affect their specific questions before selecting an approach for functional inference.

## Conclusions

Our study underscores the methodological challenges of profiling gut microbiomes from low-biomass and highly variable samples. We demonstrate that the general advantages attributed to genome-resolved metagenomics in model organisms do not necessarily apply to systems such as bats and likely many bird species that exhibit low microbial DNA content and large inter-individual variation. The relative merits of metabarcoding and metagenomics must therefore be evaluated within the specific biological and logistical context of each study, considering research objectives, resource availability, and potential biases inherent to both methods. When comparing results derived from these approaches, efforts should be made to harmonise outputs by accounting for factors that tend to overestimate diversity in metabarcoding and underestimate it in metagenomics. While ongoing initiatives such as the Earth Microbiome Project and the Earth Hologenome Initiative continue to expand microbial reference genome databases, researchers must remain aware of the technical and biological limitations that shape their data, ensuring that methodological artefacts do not obscure biological interpretation, while advocating for methodological transparency to ensure accurate cross-study comparisons.

## Code accessibility

The raw data tables containing the quantitative information and bioinformatic code scripts used for the analysis are available in a dedicated GitHub repository, both rendered into a html webbook (https://alberdilab.github.io/amplicon_shotgun_bats/) and frozen in Zenodo with Doi: XXXXXX (Aizpurua et al., 2026).

## Data accessibility

The raw 16S amplicon and metagenomic sequencing data have been deposited in the European Nucleotide Archive under Bioproject PRJEB98190.

## Acknowledgement

This project was funded through a Natural Environment Research Council Independent Research Fellowship (NE/M018660/1) awarded to OR and the Danish National Research Foundation (DNRF143) awarded to OA and AA. EJM was supported through a GW4+ NERC DTP PhD studentship and EMBO laboratory visit funding (8075). Fieldwork was also supported through a Genetic Society Travel grant and Bat Conservation International grant, both awarded to EJM.

## References

Aizpurua, O., Dunn, R. R., Hansen, L. H., Gilbert, M. T. P., & Alberdi, A. (2023). Field and laboratory guidelines for reliable bioinformatic and statistical analysis of bacterial shotgun metagenomic data. Critical Reviews in Biotechnology, 1–19.

Aizpurua, O., Martin-Bideguren, G., Gaun, N., & Alberdi, A. (2024). Dietary intervention in captive-bred hares fails to enrich gut microbiomes with wild-like functions. In bioRxiv (p. 2024.12. 03.626655). 10.1101/2024.12.03.626655

Aizpurua, O., Nyholm, L., Morris, E., Chaverri, G., Herrera Montalvo, L. G., Flores-Martinez, J. J., Lin, A., Razgour, O., Gilbert, M. T. P., & Alberdi, A. (2021). The role of the gut microbiota in the dietary niche expansion of fishing bats. Animal Microbiome, 3(1), 76.

Alberdi, A. (n.d.). distillR: R package for distilling functional annotations of bacterial genomes and metagenomes. Github. Retrieved September 25, 2023, from https://github.com/anttonalberdi/distillR

Alberdi, A., Aizpurua, O., Bohmann, K., Zepeda-Mendoza, M. L., & Gilbert, M. T. P. (2016). Do vertebrate gut metagenomes confer rapid ecological adaptation? Trends in Ecology & Evolution, 31(9), 689–699.

Alberdi, A., Aizpurua, O., Gilbert, M. T. P., & Bohmann, K. (2018). Scrutinizing key steps for reliable metabarcoding of environmental samples. Methods in Ecology and Evolution / British Ecological Society, 9, 134–147.

Alberdi, A., & Gilbert, M. T. P. (2019a). A guide to the application of Hill numbers to DNA-based diversity analyses. Molecular Ecology Resources, 19(4), 804–817.

Alberdi, A., & Gilbert, M. T. P. (2019b). hilldiv: an R package for the integral analysis of diversity based on Hill numbers. In bioRxiv (p. 545665). 10.1101/545665

Albertsen, M., Hugenholtz, P., Skarshewski, A., Nielsen, K. L., Tyson, G. W., & Nielsen, P. H. (2013). Genome sequences of rare, uncultured bacteria obtained by differential coverage binning of multiple metagenomes. Nature Biotechnology, 31(6), 533–538.

Albright, S., & Louca, S. (2023). Trait biases in microbial reference genomes. Scientific Data, 10(1), 84.

Alneberg, J., Bjarnason, B. S., de Bruijn, I., Schirmer, M., Quick, J., Ijaz, U. Z., Lahti, L., Loman, N. J., Andersson, A. F., & Quince, C. (2014). Binning metagenomic contigs by coverage and composition. Nature Methods, 11(11), 1144–1146.

Amichai, E., & Korine, C. (2020). Kuhl’s Pipistrelle Pipistrellus kuhlii (Kuhl, 1817). In Handbook of the Mammals of Europe (pp. 1–19). Springer International Publishing.

Arellano-Hernández, H. D., Montes-Carreto, L. M., Guerrero, J. A., & Martinez-Romero, E. (2024). The fecal microbiota of the mouse-eared bat (Myotis velifer) with new records of microbial taxa for bats. PloS One, 19(12), e0314847.

Aroney, S. T. N., Newell, R. J. P., Nissen, J. N., Camargo, A. P., Tyson, G. W., & Woodcroft, B. J. (2025). CoverM: read alignment statistics for metagenomics. Bioinformatics (Oxford, England), 41(4), btaf147.

Arumugam, M., Raes, J., Pelletier, E., Le Paslier, D., Yamada, T., Mende, D. R., Fernandes, G. R., Tap, J., Bruls, T., Batto, J.-M., Bertalan, M., Borruel, N., Casellas, F., Fernandez, L., Gautier, L., Hansen, T., Hattori, M., Hayashi, T., Kleerebezem, M., … Bork, P. (2011). Enterotypes of the human gut microbiome. Nature, 473(7346), 174–180.

Bateman, A., Coin, L., Durbin, R., Finn, R. D., Hollich, V., Griffiths-Jones, S., Khanna, A., Marshall, M., Moxon, S., Sonnhammer, E. L. L., Studholme, D. J., Yeats, C., & Eddy, S. R. (2004). The Pfam protein families database. Nucleic Acids Research, 32(Database issue), D138–D141.

Binladen, J., Gilbert, M. T. P., Bollback, J. P., Panitz, F., Bendixen, C., Nielsen, R., & Willerslev, E. (2007). The use of coded PCR primers enables high-throughput sequencing of multiple homolog amplification products by 454 parallel sequencing. PloS One, 2(2), e197.

Bowers, R. M., Kyrpides, N. C., Stepanauskas, R., Harmon-Smith, M., Doud, D., Reddy, T. B. K., Schulz, F., Jarett, J., Rivers, A. R., Eloe-Fadrosh, E. A., Tringe, S. G., Ivanova, N. N., Copeland, A., Clum, A., Becraft, E. D., Malmstrom, R. R., Birren, B., Podar, M., Bork, P., … Woyke, T. (2017). Minimum information about a single amplified genome (MISAG) and a metagenome-assembled genome (MIMAG) of bacteria and archaea. Nature Biotechnology, 35(8), 725–731.

Brooks, M. E., Kristensen, K., Van Benthem, K. J., Magnusson, A., Berg, C. W., Nielsen, A., Skaug, H. J., Machler, M., & Bolker, B. M. (2017). glmmTMB balances speed and flexibility among packages for zero-inflated generalized linear mixed modeling. The R Journal, 9(2), 378–400.

Callahan, B. J., McMurdie, P. J., Rosen, M. J., Han, A. W., Johnson, A. J. A., & Holmes, S. P. (2016). DADA2: High-resolution sample inference from Illumina amplicon data. Nature Methods, 13(7), 581–583.

Caporaso, J. G., Lauber, C. L., Walters, W. A., Berg-Lyons, D., Lozupone, C. A., Turnbaugh, P. J., Fierer, N., & Knight, R. (2011). Global patterns of 16S rRNA diversity at a depth of millions of sequences per sample. Proceedings of the National Academy of Sciences of the United States of America, 108 Suppl 1, 4516–4522.

Carøe, C., & Bohmann, K. (2020). Tagsteady: A metabarcoding library preparation protocol to avoid false assignment of sequences to samples. Molecular Ecology Resources, 20(6), 1620–1631.

Carøe, C., Gopalakrishnan, S., Vinner, L., Mak, S. S. T., Sinding, M.-H. S., Samaniego, J. A., Wales, N., Sicheritz-Pontén, T., & Gilbert, M. T. P. (2017). Single-tube library preparation for degraded DNA. Methods in Ecology and Evolution.

Carthey, A. J. R., Blumstein, D. T., Gallagher, R. V., Tetu, S. G., & Gillings, M. R. (2020). Conserving the holobiont. Functional Ecology, 34(4), 764–776.

Chan, P. P., & Lowe, T. M. (2019). tRNAscan-SE: Searching for tRNA Genes in Genomic Sequences. Methods in Molecular Biology, 1962, 1–14.

Chaumeil, P.-A., Mussig, A. J., Hugenholtz, P., & Parks, D. H. (2019). GTDB-Tk: a toolkit to classify genomes with the Genome Taxonomy Database. Bioinformatics, 36(6), 1925–1927.

Chen, S., Zhou, Y., Chen, Y., & Gu, J. (2018). fastp: an ultra-fast all-in-one FASTQ preprocessor. Bioinformatics, 34(17), i884–i890.

Chklovski, A., Parks, D. H., Woodcroft, B. J., & Tyson, G. W. (2022). CheckM2: a rapid, scalable and accurate tool for assessing microbial genome quality using machine learning. In bioRxiv (p. 2022.07.11.499243). 10.1101/2022.07.11.499243

Cláudio, V. C., Novaes, R. L. M., Gardner, A. L., Nogueira, M. R., Wilson, D. E., Maldonado, J. E., Oliveira, J. A., & Moratelli, R. (2023). Taxonomic re-evaluation of New World Eptesicus and Histiotus (Chiroptera: Vespertilionidae), with the description of a new genus. *Zoologia (Curitiba*, Brazil), 40(e22029), e22029.

Cornelius Ruhs, E., McFerrin, K., Jones, D. N., Cortes-Delgado, N., Ravelomanantsoa, N. A. F., Yeoman, C. J., Plowright, R. K., & Brook, C. E. (2024). Rapid GIT transit time in volant vertebrates, with implications for convergence in microbiome composition. In bioRxivorg. 10.1101/2024.08.09.607319

Dai, W., Leng, H., Li, J., Li, A., Li, Z., Zhu, Y., Li, X., Jin, L., Sun, K., & Feng, J. (2024). The role of host traits and geography in shaping the gut microbiome of insectivorous bats. mSphere, 9(4), e0008724.

Davis, N. M., Proctor, D. M., Holmes, S. P., Relman, D. A., & Callahan, B. J. (2018). Simple statistical identification and removal of contaminant sequences in marker-gene and metagenomics data. Microbiome, 6(1), 226.

DeAngelis, M. M., Wang, D. G., & Hawkins, T. L. (1995). Solid-phase reversible immobilization for the isolation of PCR products. Nucleic Acids Research, 23(22), 4742–4743.

Djemiel, C., Maron, P.-A., Terrat, S., Dequiedt, S., Cottin, A., & Ranjard, L. (2022). Inferring microbiota functions from taxonomic genes: a review. GigaScience, 11(1). 10.1093/gigascience/giab090

Douglas, G. M., Maffei, V. J., Zaneveld, J. R., Yurgel, S. N., Brown, J. R., Taylor, C. M., Huttenhower, C., & Langille, M. G. I. (2020). PICRUSt2 for prediction of metagenome functions. Nature Biotechnology, 38(6), 685–688.

Eisenhofer, R., & Alberdi, A. (2023, November 6). The Earth Hologenome Initiative Bioinformatics Workflow. https://www.earthhologenome.org/bioinformatics/

Eisenhofer, R., Alberdi, A., & Woodcroft, B. J. (2024). Large-scale estimation of bacterial and archaeal DNA prevalence in metagenomes reveals biome-specific patterns. In bioRxiv (p. 2024.05.16.594470). 10.1101/2024.05.16.594470

Eisenhofer, R., Odriozola, I., & Alberdi, A. (2023). Impact of microbial genome completeness on metagenomic functional inference. ISME Communications, 3(1), 12.

Eloe-Fadrosh, E. A., Ivanova, N. N., Woyke, T., & Kyrpides, N. C. (2016). Metagenomics uncovers gaps in amplicon-based detection of microbial diversity. Nature Microbiology, 1(4), 15032.

Feldman, R., Whitaker, J. O., & Yom-Tov, Y. (2000). Dietary composition and habitat use in a desert insectivorous bat community in Israel. Acta Chiropterologica / Museum and Institute of Zoology, Polish Academy of Sciences, 1(02). https://www.infona.pl/resource/bwmeta1.element.agro-dfe2aeaf-b164-429a-a1f0-202b37fb1c9a

Finn, R. D., Clements, J., & Eddy, S. R. (2011). HMMER web server: interactive sequence similarity searching. Nucleic Acids Research, 39(Web Server issue), W29–W37.

Flint, H. J., Scott, K. P., Louis, P., & Duncan, S. H. (2012). The role of the gut microbiota in nutrition and health. Nature Reviews. Gastroenterology & Hepatology, 9(10), 577–589.

Gibson, K. M., Nguyen, B. N., Neumann, L. M., Miller, M., Buss, P., Daniels, S., Ahn, M. J., Crandall, K. A., & Pukazhenthi, B. (2019). Gut microbiome differences between wild and captive black rhinoceros - implications for rhino health. Scientific Reports, 9(1), 7570.

Hill, M. O. (1973). Diversity and Evenness: A Unifying Notation and Its Consequences. Ecology, 54(2), 427–432.

Huerta-Cepas, J., Szklarczyk, D., Heller, D., Hernández-Plaza, A., Forslund, S. K., Cook, H., Mende, D. R., Letunic, I., Rattei, T., Jensen, L. J., von Mering, C., & Bork, P. (2019). eggNOG 5.0: a hierarchical, functionally and phylogenetically annotated orthology resource based on 5090 organisms and 2502 viruses. Nucleic Acids Research, 47(D1), D309–D314.

Iwai, S., Weinmaier, T., Schmidt, B. L., Albertson, D. G., Poloso, N. J., Dabbagh, K., & DeSantis, T. Z. (2016). Piphillin: Improved Prediction of Metagenomic Content by Direct Inference from Human Microbiomes. PloS One, 11(11), e0166104.

Jain, C., Rodriguez-R, L. M., Phillippy, A. M., Konstantinidis, K. T., & Aluru, S. (2018). High throughput ANI analysis of 90K prokaryotic genomes reveals clear species boundaries. Nature Communications, 9(1), 5114.

Johnson, J. S., Spakowicz, D. J., Hong, B.-Y., Petersen, L. M., Demkowicz, P., Chen, L., Leopold, S. R., Hanson, B. M., Agresta, H. O., Gerstein, M., Sodergren, E., & Weinstock, G. M. (2019). Evaluation of 16S rRNA gene sequencing for species and strain-level microbiome analysis. Nature Communications, 10(1), 5029.

Jones, G., Jacobs, D. S., Kunz, T. H., & Willig, M. R. (n.d.). Carpe noctem: the importance of bats as bioindicators. https://www.int-res.com/abstracts/esr/v8/n1-2/p93-115

Kandlikar, G. S., Gold, Z. J., Cowen, M. C., Meyer, R. S., Freise, A. C., Kraft, N. J. B., Moberg-Parker, J., Sprague, J., Kushner, D. J., & Curd, E. E. (2018). ranacapa: An R package and Shiny web app to explore environmental DNA data with exploratory statistics and interactive visualizations. F1000Research, 7(1734), 1734.

Kanehisa, M., & Goto, S. (2000). KEGG: kyoto encyclopedia of genes and genomes. Nucleic Acids Research, 28(1), 27–30.

Kang, D. D., Li, F., Kirton, E., Thomas, A., Egan, R., An, H., & Wang, Z. (2019). MetaBAT 2: an adaptive binning algorithm for robust and efficient genome reconstruction from metagenome assemblies. PeerJ, 7, e7359.

Korine, C. (2021). Botta’s Serotine Eptesicus bottae (Peters, 1869). 1–9.

Korine, C., & Pinshow, B. (2004). Guild structure, foraging space use, and distribution in a community of insectivorous bats in the Negev Desert. Journal of Zoology. 10.1017/S0952836903004539

Koziol, A., Odriozola, I., Leonard, A., Eisenhofer, R., San José, C., Aizpurua, O., & Alberdi, A. (2023). Mammals show distinct functional gut microbiome dynamics to identical series of environmental stressors. mBio, 14(5), e0160623.

Krijthe, J. H. (2015). Rtsne: T-Distributed Stochastic Neighbor Embedding using a Barnes-Hut Implementation. https://github.com/jkrijthe/Rtsne

Kunz, T. H., Braun de Torrez, E., Bauer, D., Lobova, T., & Fleming, T. H. (2011). Ecosystem services provided by bats: Ecosystem services provided by bats. Annals of the New York Academy of Sciences, 1223(1), 1–38.

Langille, M. G. I., Zaneveld, J., Caporaso, J. G., McDonald, D., Knights, D., Reyes, J. A., Clemente, J. C., Burkepile, D. E., Vega Thurber, R. L., Knight, R., Beiko, R. G., & Huttenhower, C. (2013). Predictive functional profiling of microbial communities using 16S rRNA marker gene sequences. Nature Biotechnology, 31(9), 814–821.

Langmead, B., & Salzberg, S. L. (2012). Fast gapped-read alignment with Bowtie 2. Nature Methods, 9(4), 357–359.

Laukens, D., Brinkman, B. M., Raes, J., De Vos, M., & Vandenabeele, P. (2016). Heterogeneity of the gut microbiome in mice: guidelines for optimizing experimental design. FEMS Microbiology Reviews, 40(1), 117–132.

Lenth, R., Buerkner, P., Herve, M., Love, J., & Riebl, H. (2020). Emmeans: Estimated marginal means, aka least-squares means, v1. 5.1. Vienna: R Core Team. https://scholar.google.dk/citations?user=JSj6m1IAAAAJ&hl=en&oi=sra

Leonard, A., Earth Hologenome Initiative Consortium, & Alberdi, A. (2024). A global initiative for ecological and evolutionary hologenomics. Trends in Ecology & Evolution, 39(7), 616–620.

Levin, D., Raab, N., Pinto, Y., Rothschild, D., Zanir, G., Godneva, A., Mellul, N., Futorian, D., Gal, D., Leviatan, S., Zeevi, D., Bachelet, I., & Segal, E. (2021). Diversity and functional landscapes in the microbiota of animals in the wild. Science, 372(6539), eabb5352.

Li, D., Liu, C.-M., Luo, R., Sadakane, K., & Lam, T.-W. (2015). MEGAHIT: an ultra-fast single-node solution for large and complex metagenomics assembly via succinct de Bruijn graph. Bioinformatics, 31(10), 1674–1676.

Li, H., Handsaker, B., Wysoker, A., Fennell, T., Ruan, J., Homer, N., Marth, G., Abecasis, G., Durbin, R., & 1000 Genome Project Data Processing Subgroup. (2009). The Sequence Alignment/Map format and SAMtools. Bioinformatics, 25(16), 2078–2079.

Lin, H., & Peddada, S. D. (2024). Multigroup analysis of compositions of microbiomes with covariate adjustments and repeated measures. Nature Methods, 21(1), 83–91.

Liu, Y.-X., Qin, Y., Chen, T., Lu, M., Qian, X., Guo, X., & Bai, Y. (2020). A practical guide to amplicon and metagenomic analysis of microbiome data. Protein & Cell, 12, 315–330.

Louca, S., Polz, M. F., Mazel, F., Albright, M. B. N., Huber, J. A., O’Connor, M. I., Ackermann, M., Hahn, A. S., Srivastava, D. S., Crowe, S. A., Doebeli, M., & Parfrey, L. W. (2018). Function and functional redundancy in microbial systems. Nature Ecology & Evolution, 2(6), 936–943.

Malmstrom, R. R., & Eloe-Fadrosh, E. A. (2019). Advancing genome-resolved metagenomics beyond the shotgun. mSystems, 4(3). 10.1128/mSystems.00118-19

Marotz, C. A., Sanders, J. G., Zuniga, C., Zaramela, L. S., Knight, R., & Zengler, K. (2018). Improving saliva shotgun metagenomics by chemical host DNA depletion. Microbiome, 6(1), 42.

Martinez Arbizu, P. (2020). pairwiseAdonis: Pairwise multilevel comparison using adonis. R package version 0.4. Github. https://github.com/pmartinezarbizu/pairwiseAdonis

Martin, M. (2011). Cutadapt removes adapter sequences from high-throughput sequencing reads. EMBnet.journal, 17(1), 10–12.

Matchado, M. S., Rühlemann, M., Reitmeier, S., Kacprowski, T., Frost, F., Haller, D., Baumbach, J., & List, M. (2024). On the limits of 16S rRNA gene-based metagenome prediction and functional profiling. Microbial Genomics, 10(2). 10.1099/mgen.0.001203

McFall-Ngai, M., Hadfield, M. G., Bosch, T. C. G., Carey, H. V., Domazet-Lošo, T., Douglas, A. E., Dubilier, N., Eberl, G., Fukami, T., Gilbert, S. F., Hentschel, U., King, N., Kjelleberg, S., Knoll, A. H., Kremer, N., Mazmanian, S. K., Metcalf, J. L., Nealson, K., Pierce, N. E., … Wernegreen, J. J. (2013). Animals in a bacterial world, a new imperative for the life sciences. Proceedings of the National Academy of Sciences of the United States of America, 110(9), 3229–3236.

McLaren, M. R., & Callahan, B. J. (2020). Pathogen resistance may be the principal evolutionary advantage provided by the microbiome. Philosophical Transactions of the Royal Society of London. Series B, Biological Sciences, 375(1808), 20190592.

Murray, D. C., Coghlan, M. L., & Bunce, M. (2015). From benchtop to desktop: important considerations when designing amplicon sequencing workflows. PloS One, 10(4), e0124671.

Muyzer, G., de Waal, E. C., & Uitterlinden, A. G. (1993). Profiling of complex microbial populations by denaturing gradient gel electrophoresis analysis of polymerase chain reaction-amplified genes coding for 16S rRNA. Applied and Environmental Microbiology, 59(3), 695–700.

Nearing, J. T., Comeau, A. M., & Langille, M. G. I. (2021). Identifying biases and their potential solutions in human microbiome studies. Microbiome, 9(1), 113.

Oksanen, J., Blanchet, F. G., Kindt, R., Legendre, P., Minchin, P. R., O’hara, R. B., Simpson, G. L., Solymos, P., Stevens, M. H. H., Wagner, H., & Others. (2013). Package “vegan.” Community Ecology Package, Version, 2(9), 1–295.

Olm, M. R., Brown, C. T., Brooks, B., & Banfield, J. F. (2017). dRep: a tool for fast and accurate genomic comparisons that enables improved genome recovery from metagenomes through de-replication. The ISME Journal, 11(12), 2864–2868.

Ondov, B. D., Treangen, T. J., Melsted, P., Mallonee, A. B., Bergman, N. H., Koren, S., & Phillippy, A. M. (2016). Mash: fast genome and metagenome distance estimation using MinHash. Genome Biology, 17(1), 132.

Oren, A., & Garrity, G. M. (2021). Valid publication of the names of forty-two phyla of prokaryotes. International Journal of Systematic and Evolutionary Microbiology, 71(10). 10.1099/ijsem.0.005056

Park, B. H., Karpinets, T. V., Syed, M. H., Leuze, M. R., & Uberbacher, E. C. (2010). CAZymes Analysis Toolkit (CAT): web service for searching and analyzing carbohydrate-active enzymes in a newly sequenced organism using CAZy database. Glycobiology, 20(12), 1574–1584.

Parks, D. H., Chuvochina, M., Chaumeil, P.-A., Rinke, C., Mussig, A. J., & Hugenholtz, P. (2020). A complete domain-to-species taxonomy for Bacteria and Archaea. Nature Biotechnology, 38(9), 1079–1086.

Parks, D. H., Chuvochina, M., Waite, D. W., Rinke, C., Skarshewski, A., Chaumeil, P.-A., & Hugenholtz, P. (2018). A standardized bacterial taxonomy based on genome phylogeny substantially revises the tree of life. Nature Biotechnology, 36(10), 996–1004.

Pereira-Marques, J., Hout, A., Ferreira, R. M., Weber, M., Pinto-Ribeiro, I., van Doorn, L.-J., Knetsch, C. W., & Figueiredo, C. (2019). Impact of host DNA and sequencing depth on the taxonomic resolution of whole metagenome sequencing for microbiome analysis. Frontiers in Microbiology, 10, 1277.

Pietroni, C., Gaun, N., Leonard, A., Lauritsen, J., Martin-Bideguren, G., Odriozola, I., Aizpurua, O., Alberdi, A., & Eisenhofer, R. (2024). Hologenomic data generation and analysis in wild vertebrates. Methods in Ecology and Evolution. 10.1111/2041-210x.14456

Poretsky, R., Rodriguez-R, L. M., Luo, C., Tsementzi, D., & Konstantinidis, K. T. (2014). Strengths and limitations of 16S rRNA gene amplicon sequencing in revealing temporal microbial community dynamics. PloS One, 9(4), e93827.

Prodan, A., Tremaroli, V., Brolin, H., Zwinderman, A. H., Nieuwdorp, M., & Levin, E. (2020). Comparing bioinformatic pipelines for microbial 16S rRNA amplicon sequencing. PloS One, 15(1), e0227434.

Quince, C., Walker, A. W., Simpson, J. T., Loman, N. J., & Segata, N. (2017). Shotgun metagenomics, from sampling to analysis. Nature Biotechnology, 35(9), 833–844.

Razgour, O., Persey, M., Shamir, U., & Korine, C. (2018). The role of climate, water and biotic interactions in shaping biodiversity patterns in arid environments across spatial scales. Diversity & Distributions, 24(10), 1440–1452.

R Core Team R, Others. (2013). R: A language and environment for statistical computing. Vienna, Austria. https://cran.r-project.org/doc/manuals/r-release/fullrefman.pdf

Rinke, C., Chuvochina, M., Mussig, A. J., Chaumeil, P.-A., Davín, A. A., Waite, D. W., Whitman, W. B., Parks, D. H., & Hugenholtz, P. (2021). A standardized archaeal taxonomy for the Genome Taxonomy Database. Nature Microbiology, 6(7), 946–959.

Rohland, N., & Reich, D. (2012). Cost-effective, high-throughput DNA sequencing libraries for multiplexed target capture. Genome Research, 22(5), 939–946.

Roswag, A., Becker, N. I., & Encarnação, J. A. (2012). Inter□ and intraspecific comparisons of retention time in insectivorous bat species (Vespertilionidae): Retention time of insectivorous bats. Journal of Zoology (London, England: 1987), 288(2), 85–92.

Roume, H., Mondot, S., Saliou, A., Le Fresne-Languille, S., & Doré, J. (2023). Multicenter evaluation of gut microbiome profiling by next-generation sequencing reveals major biases in partial-length metabarcoding approach. Scientific Reports, 13(1), 22593.

Schnell, I. B., Bohmann, K., & Gilbert, M. T. P. (2015). Tag jumps illuminated--reducing sequence-to-sample misidentifications in metabarcoding studies. Molecular Ecology Resources, 15(6), 1289–1303.

Schubert, M., Lindgreen, S., & Orlando, L. (2016). AdapterRemoval v2: rapid adapter trimming, identification, and read merging. BMC Research Notes, 9, 88.

Seemann, T., & Booth, T. (2018). Barrnap: basic rapid ribosomal RNA predictor. GitHub Repository.

Selvin, J., Lanong, S., Syiem, D., De Mandal, S., Kayang, H., Kumar, N. S., & Kiran, G. S. (2019). Culture-dependent and metagenomic analysis of lesser horseshoe bats’ gut microbiome revealing unique bacterial diversity and signatures of potential human pathogens. Microbial Pathogenesis, 137(103675), 103675.

Shaffer, M., Borton, M. A., McGivern, B. B., Zayed, A. A., La Rosa, S. L., Solden, L. M., Liu, P., Narrowe, A. B., Rodríguez-Ramos, J., Bolduc, B., Gazitúa, M. C., Daly, R. A., Smith, G. J., Vik, D. R., Pope, P. B., Sullivan, M. B., Roux, S., & Wrighton, K. C. (2020). DRAM for distilling microbial metabolism to automate the curation of microbiome function. Nucleic Acids Research, 48(16), 8883–8900.

Simmons, N. B., & Cirranello, A. L. (2020). Bat species of the world: a taxonomic and geographic database.

Steinegger, M., & Söding, J. (2017). MMseqs2 enables sensitive protein sequence searching for the analysis of massive data sets. Nature Biotechnology, 35(11), 1026–1028.

Sun, S., Jones, R. B., & Fodor, A. A. (2020). Inference-based accuracy of metagenome prediction tools varies across sample types and functional categories. Microbiome, 8(1), 46.

Tang, G., Carr, A. V., Perez, C., Ramos Sarmiento, K., Levy, L., Lampe, J. W., Diener, C., & Gibbons, S. M. (2025). Metagenomic estimation of absolute bacterial biomass in the mammalian gut through host-derived read normalization. mSystems, 10(8), e0098425.

Thomas, F., Hehemann, J.-H., Rebuffet, E., Czjzek, M., & Michel, G. (2011). Environmental and gut bacteroidetes: the food connection. Frontiers in Microbiology, 2, 93.

Uritskiy, G. V., DiRuggiero, J., & Taylor, J. (2018). MetaWRAP—a flexible pipeline for genome-resolved metagenomic data analysis. Microbiome, 6(1), 1–13.

Vengust, M., Knapic, T., & Weese, J. S. (2018). The fecal bacterial microbiota of bats; Slovenia. PloS One, 13(5), e0196728.

Větrovský, T., & Baldrian, P. (2013). The variability of the 16S rRNA gene in bacterial genomes and its consequences for bacterial community analyses. PloS One, 8(2), e57923.

Woodcroft, B. J., Aroney, S. T. N., Zhao, R., Cunningham, M., Mitchell, J. A. M., Nurdiansyah, R., Blackall, L., & Tyson, G. W. (2025). Comprehensive taxonomic identification of microbial species in metagenomic data using SingleM and Sandpiper. Nature Biotechnology, 1–6.

Worsley, S. F., Videvall, E., Harrison, X. A., Björk, J. R., Mazel, F., & Wanelik, K. M. (2024). Probing the functional significance of wild animal microbiomes using omics data. Functional Ecology. 10.1111/1365-2435.14650

Wright, R. J., Comeau, A. M., & Langille, M. G. I. (2023). From defaults to databases: parameter and database choice dramatically impact the performance of metagenomic taxonomic classification tools. Microbial Genomics, 9(3). 10.1099/mgen.0.000949

Wu, Y.-W., Simmons, B. A., & Singer, S. W. (2016). MaxBin 2.0: an automated binning algorithm to recover genomes from multiple metagenomic datasets. Bioinformatics, 32(4), 605–607.

Yom-tov, Y., & Kadmon, R. (1998). Analysis of the distribution of insectivorous bats in Israel. Diversity & Distributions, 4(2), 63–70.

Yom-Tov, Y., & Tchernov, E. (1988). The zoogeography of Israel. The distribution and abundance at a zoogeographical crossroad. cabidigitallibrary.org.

Zepeda Mendoza, M. L., Sicheritz-Pontén, T., & Gilbert, M. T. P. (2015). Environmental genes and genomes: understanding the differences and challenges in the approaches and software for their analyses. Briefings in Bioinformatics, 16(5), 745–758.

Zhao, C., Shi, Z. J., & Pollard, K. S. (2023). Pitfalls of genotyping microbial communities with rapidly growing genome collections. Cell Systems, 14(2), 160–176.e3.

