## Supplementary tables for "Impact of sequencing approach on ecological inference from highly variable bat gut microbiomes"

**Table 1.** Percentages of phyla in the three bat species recovered from (a) metagenomic and metabarcoding datasets with (b) standard filtering and (c) selected filtering.

a)

| **Phylum** | **Total** | **Cnephaeus** | **Hypsugo** | **Pipistrellus** |
| --- | --- | --- | --- | --- |
| Pseudomonadota | 68.255±37.904 | 63.683±35.64 | 89.049±29.599 | 52.374±38.209 |
| Bacillota | 17.862±28.832 | 17.716±28.111 | 5.375±19.493 | 28.713±32.41 |
| Desulfobacterota | 3.981±10.582 | 7.227±13.106 | 0±0 | 5.944±13.024 |
| Bacteroidota | 6.774±17.384 | 5.695±9.903 | 5.263±22.942 | 8.569±14.844 |
| Fusobacteriota | 0.694±1.785 | 1.818±3.061 | 0±0 | 0.781±1.589 |
| Campylobacterota | 1.288±6.836 | 1.731±2.428 | 0±0 | 2.198±10.309 |
| Elusimicrobiota | 0.141±0.755 | 0.72±1.643 | 0±0 | 0±0 |
| Synergistota | 0.405±1.282 | 0.562±1.116 | 0±0 | 0.684±1.771 |
| Planctomycetota | 0.086±0.454 | 0.44±0.985 | 0±0 | 0±0 |
| Deferribacterota | 0.08±0.4 | 0.407±0.861 | 0±0 | 0±0 |
| Actinomycetota | 0.075±0.534 | 0±0 | 0.201±0.876 | 0±0 |
| Cyanobacteriota | 0.318±1.537 | 0±0 | 0±0 | 0.737±2.302 |
| Spirochaetota | 0.041±0.296 | 0±0 | 0.111±0.486 | 0±0 |

b)

| **Phylum** | **Total** | **Cnephaeus** | **Hypsugo** | **Pipistrellus** |
| --- | --- | --- | --- | --- |
| Pseudomonadota | 58.806±25.926 | 53.499±30.532 | 67.515±18.835 | 53.697±28.06 |
| Bacillota | 26.442±20.944 | 25.488±23.748 | 22.918±16.066 | 29.919±23.604 |
| Bacteroidota | 4.876±7.757 | 5.588±8.456 | 5.332±8.486 | 4.158±7.066 |
| Fusobacteriota | 1.831±4.276 | 5.054±6.787 | 0.382±0.927 | 1.618±4.021 |
| Desulfobacterota | 2.107±4.594 | 4.449±5.799 | 0.327±0.938 | 2.58±5.419 |
| Patescibacteria | 0.48±2.558 | 2.198±5.658 | 0.052±0.151 | 0.069±0.295 |
| Rs-K70 termite group | 0.946±2.852 | 1.378±3.222 | 0.372±1.304 | 1.246±3.603 |
| Synergistota | 0.364±0.935 | 0.671±1.257 | 0.167±0.704 | 0.394±0.948 |
| Planctomycetota | 0.177±0.475 | 0.602±0.815 | 0.016±0.059 | 0.123±0.371 |
| Campylobacterota | 1.088±6.268 | 0.528±0.911 | 0.265±0.675 | 2.053±9.542 |
| Verrucomicrobiota | 0.338±1.917 | 0.137±0.347 | 0.097±0.367 | 0.638±2.901 |
| Cyanobacteriota | 1.062±4.763 | 0.119±0.289 | 0.64±2.365 | 1.856±6.924 |
| Elusimicrobiota | 0.021±0.125 | 0.105±0.278 | 0±0 | 0.002±0.01 |
| Actinomycetota | 0.783±1.852 | 0.104±0.267 | 1.624±2.793 | 0.366±0.636 |
| Deferribacterota | 0.009±0.05 | 0.044±0.111 | 0±0 | 0.002±0.005 |
| Halobacterota | 0.48±2.922 | 0.017±0.048 | 0.182±0.667 | 0.948±4.42 |
| Spirochaetota | 0.093±0.499 | 0.014±0.043 | 0.023±0.1 | 0.189±0.753 |
| Apal-E12 | 0.001±0.004 | 0.003±0.01 | 0±0 | 0±0 |
| Crenarchaeota | 0.003±0.011 | 0.001±0.002 | 0.003±0.009 | 0.003±0.014 |
| Armatimonadota | 0±0.001 | 0±0 | 0±0 | 0±0.001 |
| Bdellovibrionota | 0.006±0.028 | 0±0 | 0.005±0.015 | 0.009±0.041 |
| Chloroflexi | 0.005±0.017 | 0±0 | 0.012±0.027 | 0.002±0.006 |
| Deinococcota | 0.02±0.13 | 0±0 | 0.053±0.212 | 0±0 |
| Dependentiae | 0±0.001 | 0±0 | 0±0.001 | 0±0 |
| Euryarchaeota | 0.024±0.124 | 0±0 | 0±0 | 0.057±0.186 |
| Myxococcota | 0.004±0.021 | 0±0 | 0.011±0.035 | 0.001±0.003 |
| Nanohaloarchaeota | 0±0.002 | 0±0 | 0.001±0.003 | 0±0 |
| Sumerlaeota | 0.001±0.004 | 0±0 | 0.002±0.007 | 0±0 |
| Thermoplasmatota | 0.031±0.22 | 0±0 | 0.001±0.003 | 0.072±0.336 |
| Thermotogota | 0±0.003 | 0±0 | 0±0 | 0.001±0.004 |

| **Phylum** | **Total** | **Cnephaeus** | **Hypsugo** | **Pipistrellus** |
| --- | --- | --- | --- | --- |
| Pseudomonadota | 59.82±35.675 | 59.189±33.515 | 69.095±28.3 | 49.826±42.873 |
| Bacillota | 31.817±31.618 | 22.436±25.315 | 29.786±27.371 | 39.606±38.523 |
| Fusobacteriota | 2.473±6.759 | 5.475±8.105 | 0±0 | 3.471±8.808 |
| Bacteroidota | 1.165±5.195 | 4.116±10.646 | 0.655±2.098 | 0±0 |
| Desulfobacterota | 2.83±8.205 | 3.887±6.035 | 0±0 | 5.371±12.321 |
| Patescibacteria | 0.563±3.268 | 2.591±6.897 | 0±0 | 0±0 |
| Rs-K70 termite group | 0.9±3.222 | 1.208±3.82 | 0±0 | 1.726±4.386 |
| Synergistota | 0.15±0.713 | 0.691±1.459 | 0±0 | 0±0 |
| Campylobacterota | 0.28±1.094 | 0.406±1.283 | 0.464±1.431 | 0±0 |

**Table 2.** Summary table of bacteria found in metabarcoding standard.

| **Bat** | **TotalASVs** | **Phylum** | **Family** | **Genus** | **Lack genus** | **Lack species** | **Present in a single individual** | **Present in a single**  **species** |
| --- | --- | --- | --- | --- | --- | --- | --- | --- |
| Total | 3211 | 23 | 287 | 623 | 980 | 3027 | 2431 | 2794 |
| C. bottae | 1129 | 17 | 137 | 246 | 383 | 1075 | 693 | 857 |
| P. kuhlii | 1326 | 20 | 189 | 354 | 424 | 1251 | 874 | 975 |
| H. ariel | 1286 | 19 | 209 | 421 | 285 | 1178 | 864 | 962 |

| **Bat** | **% ASV** | **Lack genus (%)** | **Lack species (%)** | **Present in a single individual (%)** | **Present in a single species (%)** |
| --- | --- | --- | --- | --- | --- |
| Total | 100.00 | 30.52 | 94.27 | 75.71 | 87.01 |
| C. bottae | 35.16 | 33.92 | 95.22 | 61.38 | 75.91 |
| P. kuhlii | 41.30 | 31.98 | 94.34 | 65.91 | 73.53 |
| H. ariel | 40.05 | 22.16 | 91.60 | 67.19 | 74.81 |

**Table 3.** Summary table of bacteria found in metagenomics.

| **Bat** | **TotalMAGs** | **Phylum** | **Family** | **Genus** | **Lack genus** | **Lack species** | **Present in a single individual** | **Present in a single**  **species** |
| --- | --- | --- | --- | --- | --- | --- | --- | --- |
| Total | 135 | 13 | 58 | 91 | 28 | 98 | 34 | 82 |
| C. bottae | 92 | 10 | 43 | 63 | 20 | 73 | 14 | 49 |
| P. kuhlii | 69 | 8 | 38 | 54 | 12 | 44 | 11 | 19 |
| H. ariel | 30 | 5 | 17 | 24 | 5 | 13 | 9 | 14 |

| **Bat** | **% MAGs** | **Lack genus** | **Lack species (%)** | **Present in a single individual (%)** | **Present in a single species (%)** |
| --- | --- | --- | --- | --- | --- |
| Total | 100.00 | 20.74 | 72.59 | 25.19 | 60.74 |
| C. bottae | 68.15 | 21.74 | 79.35 | 15.22 | 53.26 |
| P. kuhlii | 51.11 | 17.39 | 63.77 | 15.94 | 27.54 |
| H. ariel | 22.22 | 16.67 | 43.33 | 30.00 | 46.67 |

**Table 4.** Summary table of bacteria found in metabarcoding selected.

| **Bat** | **TotalASVs** | **Phylum** | **Family** | **Genus** | **Lack genus** | **Lack species** | **Present in a single individual** | **Present in a single**  **species** |
| --- | --- | --- | --- | --- | --- | --- | --- | --- |
| Total | 165 | 9 | 51 | 71 | 37 | 150 | 96 | 141 |
| C. bottae | 130 | 9 | 40 | 55 | 29 | 118 | 96 | 108 |
| P. kuhlii | 19 | 5 | 11 | 12 | 2 | 17 | 0 | 1 |
| H. ariel | 45 | 4 | 25 | 28 | 9 | 42 | 0 | 32 |

| **Bat** | **% ASV** | **Lack genus (%)** | **Lack species (%)** | **Present in a single individual (%)** | **Present in a single species (%)** |
| --- | --- | --- | --- | --- | --- |
| Total | 100.00 | 22.42 | 90.91 | 58.18 | 85.45 |
| C. bottae | 78.79 | 22.31 | 90.77 | 73.85 | 83.08 |
| P. kuhlii | 11.52 | 10.53 | 89.47 | 0.00 | 5.26 |
| H. ariel | 27.27 | 20.00 | 93.33 | 0.00 | 71.11 |

**Table 5.** Alpha diversity values at a) ASV or MAG level, b) family level and c) phylum level.

a)

| **Approach** | **Metric** | **Total** | **Cnephaeus** | **Pipistrellus** | **Hypsugo** |
| --- | --- | --- | --- | --- | --- |
| Metagenomics | Richness | 9±11.45 | 19.7±18.6 | 8.82±8.73 | 3.58±2.73 |
| Metagenomics | Neutral | 4.42±5.06 | 7.33±6.87 | 5.05±5.48 | 2.15±1.42 |
| Metabarcoding Standard filtering | Richness | 118.45±111.27 | 185.7±181.29 | 97.91±99.87 | 106.84±56.14 |
| Metabarcoding Standard filtering | Neutral | 20.64±20.61 | 31.16±35.42 | 17.99±17.71 | 18.16±10.51 |
| Metabarcoding  Selected filtering | Richness | 7.87±6.17 | 15.1±6.23 | 4.12±3.69 | 7.42±4.6 |
| Metabarcoding  Selected filtering | Neutral | 5.52±4.44 | 9.69±4.66 | 3.14±2.66 | 5.46±4.14 |

b)

| **Approach** | **Metric** | **Total** | **Cnephaeus** | **Pipistrellus** | **Hypsugo** |
| --- | --- | --- | --- | --- | --- |
| Metagenomics | richness | 6.47±6.67 | 12.2±10.51 | 6.73±5.35 | 3.16±2.27 |
| Metagenomics | neutral | 3.2±2.79 | 4.63±4.12 | 3.64±2.75 | 1.94±1.12 |
| Metabarcoding Standard filtering | richness | 38.61±20.88 | 40.4±20.86 | 33.73±22.76 | 43.32±18.28 |
| Metabarcoding Standard filtering | neutral | 7.93±5.03 | 8.65±6.76 | 6.74±5.19 | 8.94±3.61 |
| Metabarcoding  Selected filtering | richness | 4.96±3.53 | 7.8±4.29 | 2.65±1.84 | 5.53±3.03 |
| Metabarcoding  Selected filtering | neutral | 3.45±2.4 | 4.58±2.73 | 2.11±1.35 | 4.06±2.5 |

c)

| **Approach** | **Metric** | **Total** | **Cnephaeus** | **Pipistrellus** | **Hypsugo** |
| --- | --- | --- | --- | --- | --- |
| Metagenomics | richness | 2.78±2.21 | 5.1±2.96 | 3.09±1.66 | 1.21±0.54 |
| Metagenomics | neutral | 1.77±1.11 | 2.37±1.43 | 2.09±1.14 | 1.07±0.2 |
| Metabarcoding Standard filtering | richness | 8.04±3.65 | 10±4.16 | 7.27±3.83 | 7.89±2.9 |
| Metabarcoding Standard filtering | neutral | 2.55±1.08 | 2.98±1.37 | 2.58±1.19 | 2.28±0.69 |
| Metabarcoding  Selected filtering | richness | 2.37±1.39 | 3.6±1.84 | 2.12±1.32 | 1.95±0.71 |
| Metabarcoding  Selected filtering | neutral | 1.82±0.79 | 2.3±0.9 | 1.74±0.91 | 1.63±0.5 |

Table 6. PERMANOVA results of beta diversities differences between metagenomics and metabarcoding with standard filtering at a) the family level and b) phylum level, and metagenomics and metabarcoding with selected filtering at c) the family level and d) phylum level.

a)

q=0

| **term** | **df** | **SumOfSqs** | **R2** | **statistic** | **p.value** |
| --- | --- | --- | --- | --- | --- |
| Species | 2 | 2.129351 | 0.05344404 | 3.159414 | 0.001 |
| method | 1 | 4.660736 | 0.11697862 | 13.830689 | 0.001 |
| Species:method | 2 | 1.375945 | 0.03453450 | 2.041552 | 0.002 |
| Residual | 94 | 31.676598 | 0.79504284 | NA | NA |
| Total | 99 | 39.842630 | 1.00000000 | NA | NA |

q=1

| **term** | **df** | **SumOfSqs** | **R2** | **statistic** | **p.value** |
| --- | --- | --- | --- | --- | --- |
| Species | 2 | 2.428541 | 0.06100168 | 3.352047 | 0.001 |
| method | 1 | 2.202473 | 0.05532317 | 6.080026 | 0.001 |
| Species:method | 2 | 0.766532 | 0.01925426 | 1.058023 | 0.413 |
| Residual | 95 | 34.413497 | 0.86442089 | NA | NA |
| Total | 100 | 39.811043 | 1.00000000 | NA | NA |

b)

q=0

| **term** | **df** | **SumOfSqs** | **R2** | **statistic** | **p.value** |
| --- | --- | --- | --- | --- | --- |
| Species | 2 | 2.343946 | 0.10341297 | 8.051068 | 0.001 |
| method | 1 | 5.002004 | 0.22068430 | 34.362115 | 0.001 |
| Species:method | 2 | 1.636595 | 0.07220521 | 5.621432 | 0.001 |
| Residual | 94 | 13.683335 | 0.60369752 | NA | NA |
| Total | 99 | 22.665879 | 1.00000000 | NA | NA |

q=1

| **term** | **df** | **SumOfSqs** | **R2** | **statistic** | **p.value** |
| --- | --- | --- | --- | --- | --- |
| Species | 2 | 1.0276449 | 0.10265577 | 5.662338 | 0.003 |
| method | 1 | 0.1687487 | 0.01685702 | 1.859615 | 0.156 |
| Species:method | 2 | 0.1935300 | 0.01933252 | 1.066353 | 0.382 |
| Residual | 95 | 8.6206674 | 0.86115470 | NA | NA |
| Total | 100 | 10.0105910 | 1.00000000 | NA | NA |

c)

q=0

| **term** | **df** | **SumOfSqs** | **R2** | **statistic** | **p.value** |
| --- | --- | --- | --- | --- | --- |
| Species | 2 | 2.648151 | 0.06670721 | 3.543779 | 0.001 |
| method | 1 | 2.517837 | 0.06342460 | 6.738783 | 0.001 |
| Species:method | 2 | 1.278723 | 0.03221118 | 1.711199 | 0.010 |
| Residual | 89 | 33.253404 | 0.83765701 | NA | NA |
| Total | 94 | 39.698115 | 1.00000000 | NA | NA |

q=1

| **term** | **df** | **SumOfSqs** | **R2** | **statistic** | **p.value** |
| --- | --- | --- | --- | --- | --- |
| Species | 2 | 2.8205988 | 0.07131605 | 3.7398991 | 0.001 |
| method | 1 | 2.0914729 | 0.05288083 | 5.5462676 | 0.001 |
| Species:method | 2 | 0.7000129 | 0.01769913 | 0.9281638 | 0.533 |
| Residual | 90 | 33.9386014 | 0.85810399 | NA | NA |
| Total | 95 | 39.5506860 | 1.00000000 | NA | NA |

d)

q=0

| **term** | **df** | **SumOfSqs** | **R2** | **statistic** | **p.value** |
| --- | --- | --- | --- | --- | --- |
| Species | 2 | 2.7746936 | 0.16394000 | 10.224515 | 0.001 |
| method | 1 | 1.0793787 | 0.06377401 | 7.954842 | 0.002 |
| Species:method | 2 | 0.9947267 | 0.05877244 | 3.665485 | 0.002 |
| Residual | 89 | 12.0762564 | 0.71351355 | NA | NA |
| Total | 94 | 16.9250555 | 1.00000000 | NA | NA |

q=1

| **term** | **df** | **SumOfSqs** | **R2** | **statistic** | **p.value** |
| --- | --- | --- | --- | --- | --- |
| Species | 2 | 1.29254282 | 0.10270070 | 5.4836017 | 0.002 |
| method | 1 | 0.62148101 | 0.04938060 | 5.2732556 | 0.018 |
| Species:method | 2 | 0.06453168 | 0.00512745 | 0.2737751 | 0.849 |
| Residual | 90 | 10.60697513 | 0.84279125 | NA | NA |
| Total | 95 | 12.58553064 | 1.00000000 | NA | NA |

Table 7. ANCOMBC analysis results when using a) metabarcoding standard approach and b) metagenomics

a)

| **Phylum** | **Lfc** | **Adjusted p** | **Comparison** | **Enrichment** |
| --- | --- | --- | --- | --- |
| Cyanobacteriota | -1.6068 | 0.0442 | H. ariel vs C. bottae | Enriched in C. bottae |
| Synergistota | -2.1441 | 0.0127 | H. ariel vs C. bottae | Enriched in C. bottae |
| Actinomycetota | 1.9550 | 0.0292 | H. ariel vs C. bottae | Enriched in H. ariel |
| Planctomycetota | -2.0717 | 0.0420 | H. ariel vs C. bottae | Enriched in C. bottae |
| Patescibacteria | -2.1132 | 0.0330 | H. ariel vs C. bottae | Enriched in C. bottae |
| Patescibacteria | -2.0409 | 0.0335 | P. kuhlii vs C. bottae | Enriched in C. bottae |
| Planctomycetota | 2.0683 | 0.0375 | P. kuhlii vs H. ariel | Enriched in P. kuhlii |

b)

| **Phylum** | **Lfc** | **Adjusted p** | **Comparison** | **Enrichment** |
| --- | --- | --- | --- | --- |
| Bacteroidota | -2.4695 | 0.0159 | H. ariel vs C. bottae | Enriched in C. bottae |
| Bacteroidota | 2.5999 | 0.0159 | P. kuhlii vs H. ariel | Enriched in P. kuhlii |

Table 8. Pairwise comparison in each methodological approach at a) family and b) phylum level using the two beta diversity metrics

a) **q=0**,

Metagenomics

| **pairs** | **Df** | **SumsOfSqs** | **F.Model** | **R2** | **p.value** | **p.adjusted** | **sig** |
| --- | --- | --- | --- | --- | --- | --- | --- |
| Cb vs Ha | 1 | 0.9883724 | 2.634791 | 0.09201362 | 0.005 | 0.015 | . |
| Cb vs Pk | 1 | 0.7629461 | 2.012157 | 0.06285604 | 0.027 | 0.081 |  |
| Ha vs Pk | 1 | 1.0886497 | 2.834542 | 0.06941531 | 0.002 | 0.006 | * |

Metabarcoding standard

| **pairs** | **Df** | **SumsOfSqs** | **F.Model** | **R2** | **p.value** | **p.adjusted** | **sig** |
| --- | --- | --- | --- | --- | --- | --- | --- |
| Cb vs Ha | 1 | 1.0497877 | 3.756659 | 0.12214132 | 0.001 | 0.003 | * |
| Cb vs Pk | 1 | 0.5165122 | 1.558997 | 0.04939946 | 0.042 | 0.126 |  |
| Ha vs Pk | 1 | 0.7140756 | 2.332190 | 0.05642551 | 0.001 | 0.003 | * |

Metabarcoding selected

| **pairs** | **Df** | **SumsOfSqs** | **F.Model** | **R2** | **p.value** | **p.adjusted** | **sig** |
| --- | --- | --- | --- | --- | --- | --- | --- |
| Cb vs Ha | 1 | 1.1892164 | 3.212674 | 0.10633532 | 0.001 | 0.003 | * |
| Cb vs Pk | 1 | 0.7483175 | 1.901197 | 0.07067333 | 0.038 | 0.114 |  |
| Ha vs Pk | 1 | 0.9556233 | 2.549407 | 0.06975234 | 0.004 | 0.012 | . |

**q=1**

***Metagenomics***

| **pairs** | **Df** | **SumsOfSqs** | **F.Model** | **R2** | **p.value** | **p.adjusted** | **sig** |
| --- | --- | --- | --- | --- | --- | --- | --- |
| Cb vs Ha | 1 | 0.6990673 | 1.872374 | 0.06717669 | 0.060 | 0.180 |  |
| Cb vs Pk | 1 | 0.7388943 | 1.957133 | 0.06124244 | 0.053 | 0.159 |  |
| Ha vs Pk | 1 | 1.0166594 | 2.543328 | 0.06273112 | 0.005 | 0.015 | . |

***Metabarcoding standard***

| **pairs** | **Df** | **SumsOfSqs** | **F.Model** | **R2** | **p.value** | **p.adjusted** | **sig** |
| --- | --- | --- | --- | --- | --- | --- | --- |
| Cb vs Ha | 1 | 1.2490797 | 4.112390 | 0.13217854 | 0.001 | 0.003 | * |
| Cb vs Pk | 1 | 0.5872306 | 1.628530 | 0.05148927 | 0.080 | 0.240 |  |
| Ha vs Pk | 1 | 0.5341731 | 1.530168 | 0.03775379 | 0.092 | 0.276 |  |

***Metabarcoding selected***

| **pairs** | **Df** | **SumsOfSqs** | **F.Model** | **R2** | **p.value** | **p.adjusted** | **sig** |
| --- | --- | --- | --- | --- | --- | --- | --- |
| Cb vs Ha | 1 | 1.3532821 | 3.840951 | 0.12454063 | 0.001 | 0.003 | * |
| Cb vs Pk | 1 | 0.7636118 | 2.006946 | 0.07431220 | 0.043 | 0.129 |  |
| Ha vs Pk | 1 | 0.7412715 | 1.995708 | 0.05544294 | 0.047 | 0.141 |  |

b) **q=0**,

***Metagenomics***

| **pairs** | **Df** | **SumsOfSqs** | **F.Model** | **R2** | **p.value** | **p.adjusted** | **sig** |
| --- | --- | --- | --- | --- | --- | --- | --- |
| Cb vs Ha | 1 | 1.6935455 | 12.450959 | 0.3238140 | 0.001 | 0.003 | * |
| Cb vs Pk | 1 | 0.6958001 | 4.160066 | 0.1217815 | 0.005 | 0.015 | . |
| Ha vs Pk | 1 | 1.9319002 | 14.790370 | 0.2801717 | 0.001 | 0.003 | * |

***Metabarcoding standard***

| **pairs** | **Df** | **SumsOfSqs** | **F.Model** | **R2** | **p.value** | **p.adjusted** | **sig** |
| --- | --- | --- | --- | --- | --- | --- | --- |
| Cb vs Ha | 1 | 0.6086184 | 4.449912 | 0.14149204 | 0.001 | 0.003 | * |
| Cb vs Pk | 1 | 0.4967691 | 2.856505 | 0.08693879 | 0.020 | 0.060 |  |
| Ha vs Pk | 1 | 0.2991202 | 1.732330 | 0.04252960 | 0.103 | 0.309 |  |

***Metabarcoding selected***

| **pairs** | **Df** | **SumsOfSqs** | **F.Model** | **R2** | **p.value** | **p.adjusted** | **sig** |
| --- | --- | --- | --- | --- | --- | --- | --- |
| Cb vs Ha | 1 | 0.4237598 | 3.707872 | 0.12074662 | 0.012 | 0.036 | . |
| Cb vs Pk | 1 | 0.2966148 | 1.614777 | 0.06067220 | 0.170 | 0.510 |  |
| Ha vs Pk | 1 | 0.4132832 | 2.956948 | 0.08001061 | 0.048 | 0.144 |  |

**q=1**

***Metagenomics***

| **pairs** | **Df** | **SumsOfSqs** | **F.Model** | **R2** | **p.value** | **p.adjusted** | **sig** |
| --- | --- | --- | --- | --- | --- | --- | --- |
| Cb vs Ha | 1 | 0.2749467 | 2.822351 | 0.09792232 | 0.074 | 0.222 |  |
| Cb vs Pk | 1 | 0.1592735 | 1.095082 | 0.03521720 | 0.307 | 0.921 |  |
| Ha vs Pk | 1 | 0.9744265 | 7.700745 | 0.16850371 | 0.001 | 0.003 | * |

***Metabarcoding standard***

| **pairs** | **Df** | **SumsOfSqs** | **F.Model** | **R2** | **p.value** | **p.adjusted** | **sig** |
| --- | --- | --- | --- | --- | --- | --- | --- |
| Cb vs Ha | 1 | 1.2490797 | 4.112390 | 0.13217854 | 0.001 | 0.003 | * |
| Cb vs Pk | 1 | 0.5872306 | 1.628530 | 0.05148927 | 0.088 | 0.264 |  |
| Ha vs Pk | 1 | 0.5341731 | 1.530168 | 0.03775379 | 0.104 | 0.312 |  |

***Metabarcoding selected***

| **pairs** | **Df** | **SumsOfSqs** | **F.Model** | **R2** | **p.value** | **p.adjusted** | **sig** |
| --- | --- | --- | --- | --- | --- | --- | --- |
| Cb vs Ha | 1 | 0.1124825 | 1.557418 | 0.05453638 | 0.249 | 0.747 |  |
| Cb vs Pk | 1 | 0.1735256 | 1.121532 | 0.04293514 | 0.327 | 0.981 |  |
| Ha vs Pk | 1 | 0.3069925 | 2.827677 | 0.07678131 | 0.117 | 0.351 |  |
